# Impact of Alzheimer’s disease risk factors and local neuromelanin content on the transcriptomic landscape of the human locus coeruleus

**DOI:** 10.1101/2025.10.29.685354

**Authors:** Bernard Mulvey, Heena R. Divecha, Madhavi Tippani, Svitlana V. Bach, Rahul Bharadwaj, Ishbel Del Rosario, Sarah E. Maguire, Ryan A. Miller, Aaron J. Salisbury, Atharv Chandra, Beau A. Oster, Kelsey D. Montgomery, Sang Ho Kwon, Haya A. Algrain, Alexis R. Papariello, Louise A. Huuki-Myers, Joel E. Kleinman, Leonardo Collado-Torres, Thomas M. Hyde, Shizhong Han, Stephanie C. Hicks, Daniel R. Weinberger, Stephanie C. Page, Kristen R. Maynard, Keri Martinowich

## Abstract

The locus coeruleus (LC) is a small noradrenergic nucleus in the dorsal pons that sends projections across the brain regulating sleep, arousal, attention, stress responses, and some forms of cognition. LC neurons show pathology in the earliest stages of Alzheimer’s disease (AD), including age-related accumulation of hyperphosphorylated tau (pTau) and accelerated loss of neuromelanin (NM) pigmentation. NM-sensitive neuroimaging of the LC predicts previous cognitive decline, clinical severity, and future AD progression. While these findings suggest that the LC plays an etiologic role in AD, the molecular landscape of the LC prior to clinical manifestation of sporadic AD remains largely uncharacterized. This information is critical for developing interventions that preserve LC integrity and function. We performed spatially-resolved transcriptomics on 85 sections of human postmortem LC from *N*=33 neurotypical middle-aged donors, balanced for epidemiologic AD risk factors including sex, African or European ancestry, and *APOE* genotype (carriers of the E4/risk or E2/protective alleles). Comparing across *APOE* genotypes, we find astrocytic gene expression differences proximal to LC neurons. Associating NM content with local gene expression, we show that higher overall *APOE* gene expression correlates with reduced NM content and an enrichment of NM-associated genes in aging pathways. Unexpectedly, we find enriched LC expression of cholesterol synthesis genes, alongside evidence for lipid synthesis gene regulatory network activity in NM-containing LC specifically, revealing a potential intersection between intrinsic lipid metabolism in LC neurons, NM, and the role of APOE-mediated lipid biology in AD. Together, these data illuminate the molecular features of the human LC at spatial resolution with unprecedented sampling depth, revealing how AD risk factors and NM content influence resilience and susceptibility of this critical brain nucleus to pathology accumulation and degeneration.

## 1 | Introduction

Alzheimer’s disease (AD) is a progressive neurodegenerative disorder and the leading cause of dementia in older adults. Despite decades of research, the cellular and molecular events that initiate accumulation of pathology and AD onset are not fully understood. One of the earliest sites of AD pathology is the locus coeruleus (LC)^1^, a small brainstem nucleus that serves as the brain’s primary source of norepinephrine (NE). Evidence of soluble hyperphosphorylated tau (pTau), the precursor to neurofibrillary tangles, has been observed in the LC decades before—and regardless of—clinical AD symptom onset. Early and pre-AD stages are characterized by LC hyperexcitability^2–7^ and loss of LC axons in target brain regions^1,8–12^, likely impacting key roles of NE signaling in cognition^13–22^, sleep^23–25^, intraparenchymal brain vascular permeability^26–28^, and regulation of glial activities^9,29–34^, including inflammatory responses to AD pathology^35–37^. pTau burden in the LC predicts AD severity^3,7,38–45^, and physical propagation of pTau along forebrain-projecting axons of LC neurons may spread pTau pathology and increase vulnerability of forebrain regions to neurodegeneration^10,40,46,47^.

Despite its established and early role in AD pathogenesis, the human LC remains poorly characterized at the molecular level, largely due to technical issues related to its small size and deep location in the brain stem. Moreover, tissue studies of the LC in the context of AD have generally focused on advanced disease, while investigations of the at-risk state before clinical illness are lacking. LC-NE neurons are characterized by the presence of the pigment neuromelanin (NM), an iron-laden^48–50^ byproduct of catecholamine synthesis and metabolism that accumulates progressively through at least middle age^51^. Loss of NM-rich catecholamine neurons is observed in multiple neurodegenerative diseases, including LC neurons in AD^1,52^ and both dopaminergic substantia nigra neurons and noradrenergic LC neurons in Parkinson’s disease (PD)^53,54^. While the mechanisms by which NM contributes to cellular vulnerability in these disorders remain unclear, dual hypotheses have suggested both protective and destructive roles of NM. In early phases of AD, sequestration of oxidative catecholamine metabolites, facilitation of metal chelation, and radical scavenging by NM may be protective^49,55^. However, extracellular release of NM and its neurotoxic contents following neuronal injury or death in later disease states may activate microglia and drive inflammation^56,57^. NM bodies molecularly and structurally resemble lysosomes^56^—degradative organelles, which receive autophagosomes containing cellular debris, misfolded proteins, and pTau^58^—further suggesting that common pathways may underlie NM production and pTau clearance. Moreover, the high iron content of NM allows for measurement by specialized MRI sequences, which has demonstrated that NM loss is associated with brain-wide pathologic burden and symptom severity in AD^40^. Although cell death is the primary mechanism for loss of NM during normal aging and in neurodegenerative disease, evidence from humans^39,59^ and transgenic NM-producing mice^60^ suggests that NM can also be actively cleared; therefore, understanding the molecular correlates of NM homeostasis in intact LC neurons may provide insights into early mechanisms of neurodegenerative pathogenesis and subsequent LC loss. Finally, spatial heterogeneity in human LC cell density, cytoarchitecture, pTau accumulation, and NM loss may all play roles in spatially heterogeneous susceptibility of the LC to neurodegeneration^52,61–64^. Therefore, a more refined and spatially-oriented understanding of molecular complexity—including in relation to NM—in the LC is essential for identifying early targets of pathology and pathways that may underlie vulnerability and resilience.

A number of factors contribute to epidemiological risk for late-onset AD. First, women exhibit a higher lifetime risk of developing AD^65^, with greater impacts of pathology on cognition^66^ and brain structure^67^. Second, individuals with African ancestry (AA) are at greater overall risk relative to those with European ancestry (EA)^68–71^. Third, variation in apolipoprotein E (*APOE*) haplotype is the strongest common (allele frequency>5%) heritable risk factor for late-onset AD. Individuals carrying the *APOE4* (E4) allele have significantly increased risk, especially in the case of homozygosity (E4/E4), whereas carriers of *APOE2* (E2) allele(s) are at reduced risk^69,72^, indicating a central role in molecular susceptibility/resilience to AD. Notably, these genotype–phenotype associations vary across ancestry: E4 is strongly associated with AD risk in EA individuals, while among AA individuals, E4-associated risk is markedly attenuated^68,69,73^. These ancestry-dependent effects suggest that genomic context may modify the biological impact of *APOE* variants, providing an opportunity to explore both vulnerability and resilience mechanisms in human brain tissue. Finally, data from mice indicates that human E4 expression increases LC tau pathology and cytosolic content of toxic NE metabolites via E4 disruption of vesicular monoamine transporter 2 (VMAT2) function^74^, indicating a direct mechanism for E4-mediated LC susceptibility to AD.

To investigate how major biological risk factors for AD (*APOE* haplotype, genomic ancestry, and sex) impact molecular organization in the human LC and whether these factors impact surrounding cell populations, we generated spatially-resolved transcriptomics (SRT) data (10x Genomics Visium) from middle-aged neurotypical brain donors stratified across these risk factors. These data reveal gene expression changes associated with sex, *APOE* haplotype, genomic ancestry, and NM pigmentation. Comparison of LC transcriptomes in *APOE* E4 and E2 carriers unexpectedly shows that the strongest differences in gene expression involve canonical astrocyte genes, suggesting that *APOE* haplotype influences the density and/or transcriptional state of LC-localized astrocytes. We further show that while *APOE* haplotype effects on pontine astrocytes are shared across ancestries, E4 effects on the LC are largely ancestry-specific and more profound in EA individuals. Transcriptomic comparisons also show widespread sex differences in genes related to metabolic pathways, including male-biased expression of genes involved in oxidative phosphorylation and cholesterol synthesis. Finally, leveraging high-resolution SRT histological images, we stratify LC into NM-rich and -poor zones and demonstrate that *APOE* expression is associated with reduced NM content, while genes involved in catecholamine transmission and autophagy are associated with greater NM content. Our unique study design enabled examination of how molecular features associated with AD risk and resilience vary by haplotype, ancestry, sex, and NM within the intact spatial and cellular architecture of the LC and surrounding dorsal pons, providing a framework for understanding early mechanisms of vulnerability and resilience to neurodegeneration and data that may guide efforts to preserve LC function in aging and potentially delay or prevent the emergence of clinical AD.

## 2 | Results

### 2.1 | Transcriptional niches of the middle-aged human dorsal pons

We profiled gene expression in the human LC by generating spatially-resolved transcriptomics (SRT) data with the 10x Genomics Visium platform to investigate the molecular impact of AD risk factors in the human brain. The experimental design included *N*=33 middle-aged neurotypical brain donors with no history of clinical AD symptoms, carrying either the AD-protective E2 allele (*n*=12) or the AD-risk-conferring E4 allele of *APOE* (*n*=21). Donors were further stratified across additional AD risk factors, including biological sex and predominant global genomic ancestry (*n*=13 EA and *n*=20 AA; *Methods*) (**Fig. 1A, Table S1**). Fresh frozen tissue blocks were dissected at the level of the dorsal pons. Presence of the LC was assessed by visualizing NM using Hematoxylin and Eosin (H&E) staining and LC marker gene expression (*TH*, *DBH*) with single molecule fluorescence *in situ* hybridization (smFISH, **Fig. S1**). For neuroanatomical orientation, we also probed for serotonergic (5-HT) neurons of the raphe nucleus (*TPH2*) and GABAergic neurons (*GAD2*) of pericoerulear areas (peri-LC). Following anatomical validation, tissue blocks were scored to isolate the LC. For each donor, two consecutive cryosections were mounted per capture area with the exception of Br6263 (**Fig. 1A-B**). Tissue replicates were prepared for a subset of donors (*n*=10/33) by mounting two additional tissue sections onto a second capture area, collected from either the same tissue block (*n*=6/33) or a tissue block from the opposite hemisphere of the same donor (*n*=4/33, **Table S1**). In total, we analyzed 43 capture areas that we divided *in silico* to process each tissue section as an individual sample (85 tissue sections, referred to as “samples” throughout). Data preprocessing removed local outlier SRT spots^75^ and spots with especially low gene or UMI content, resulting in a final set of 132,484 spots containing a median of 1,202 total UMIs and 566 unique genes per spot (**Fig. S2**). Large deviations in these metrics per spot reflected the stark differences in RNA content between large, widely-projecting neurons in the LC and surrounding white matter (WM) tracts (**Table S1**). Consistent with smFISH, a dense area of gene expression corresponding to LC markers was present in most samples (**Fig. 1B**). For each section, we segmented NM pixels from high-resolution H&E images (**Fig. 1C**, **Fig. S3**) and registered these to gene expression spot coordinates to annotate spot-level NM content. Analysis of combined image and gene expression SRT data was used to identify associations between NM content and local gene expression (**Fig. 1D**).

**Figure 1.**
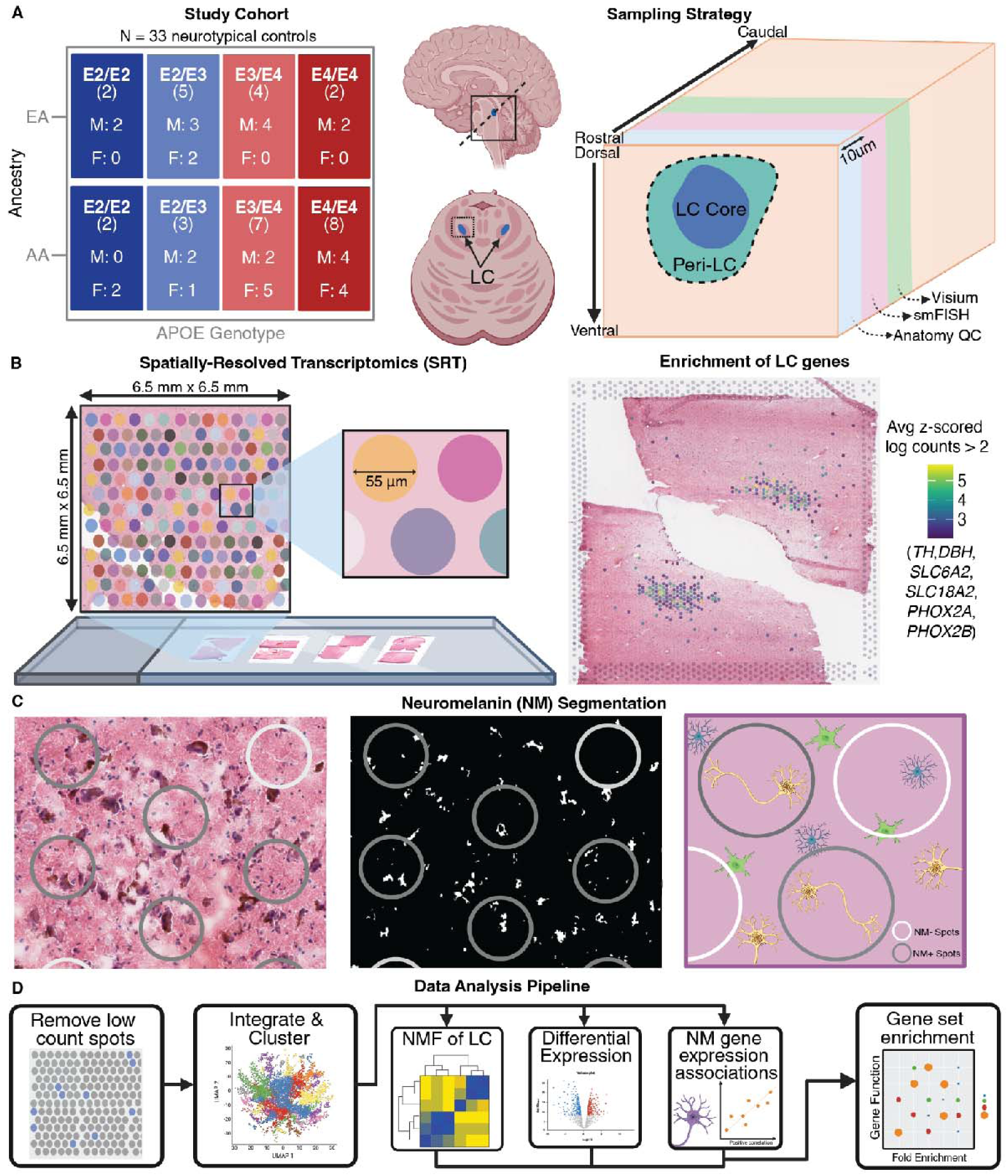
Overview of study cohort, data generation, and analyses. **A)** The cohort included 33 middle-age neurotypical human brain donors with no clinical symptoms of AD that were stratified across the *APOE* AD-protective E2 allele or the AD-risk-conferring E4 allele. Donors were further stratified across biological sex and predominant genomic (global) ancestry (EA or AA). Tissue blocks were dissected from transverse slabs of the brainstem at the level of the pons. Presence of LC in tissue blocks was confirmed by visualizing: 1) Neuromelanin (NM) on H&E stained tissues sections and 2) LC marker genes (*TH*, *DBH*) using single molecule fluorescence *in situ* hybridization (smFISH) on adjacent tissue sections. Diagram indicates sampling strategy. **B)** Additional scored tissue sections were then mounted for SRT. Briefly, tissue is mounted onto slides containing barcoded, poly-dT probes to allow for RNA capture with incorporation of a sequence corresponding to a spatial coordinate on the slide. H&E staining and imaging are performed and images are aligned to the Visium capture area, allowing visualization, image processing, and analysis of joint histology and transcriptomics data. An example SRT sample illustrating expression of LC marker genes from two consecutive tissue sections from a single donor (Br1105). **C)** NM was segmented from H&E images for incorporation into downstream analyses. *k*-means color based segmentation of the H&E image (left) was used to separate NM signal from H&E stains; the isolated NM color was then refined into binary segmentations (middle) with thresholding and artifact removal using morphological operations (**Fig. S3**). Image registration facilitated later annotation of spot-level NM content (pixel intensity and proportion of spot area with NM) and identification of LC^NM+^ and LC^NM-^ spots (right). **D**) Following quality control (QC) and clustering, analysis focused on: 1) non-negative matrix factorization (NMF) to identify transcriptionally distinct groups of LC spots; 2) differential expression (DE) analyses for each cell type and AD risk factor (sex, ancestry, *APOE* haplotype); 3) association of spot-level NM characteristics with spot-level gene expression using linear mixed models (LMMs). Gene set enrichment analyses (GSEA) was used to functionally annotate LC transcriptomic populations, genes DE across high- and low-risk AD groups for each risk factor in each domain, and to identify genes associated with NM.

We first clustered the transcriptomic data from all *N*=33 donors, focusing on annotating clusters corresponding to LC. Low- and high-stringency gene expression definitions of LC were assigned (*Methods*) to benchmark annotation accuracy using different clustering parameters. We identified feature sets comprised of highly variable genes (HVGs), high-deviance genes (HDGs)^76^, spatially-variable genes (SVGs)^77^ found in ≥43 (*i.e.*, >50% of) samples (tissue sections), and combinations of these feature sets (*e.g.*, the union of top HVGs and top SVGs; **Table S2**). Dimensionality reduction and sample-level batch correction were performed using *Harmony*^78^, followed by Louvain clustering^79^ at multiple resolutions for each feature set. Regardless of feature set or clustering granularity, a single LC cluster generally emerged in each sample. The resulting LC cluster(s) were then assessed for consistency with manual labels assigned based solely on expression of canonical LC gene markers. We selected a combination of overlapping SVGs and HDGs as the features with which to perform PCA, batch correction, and Louvain clustering for final ‘domain’ annotations. PCA and *Harmony* plots confirmed that integration was successful (**Fig. S5**, **Fig. S4**, **Fig. S6**), with highly consistent transcriptomes between the resulting LC cluster and expression-defined LC spots in nearly all donors (**Fig. S7**). Altogether, clustering yielded 9 ‘domains’ (LC, Astro, Astro-Oligo, Oligo, Oligo-Astro, Oligo-Other Cell Types Mixed, Vascular, and two domains with shared markers *ACHE* and *SLC6A4*) (**Fig. 2A**, **Table S3**, **Data S1**).

**Figure 2.**
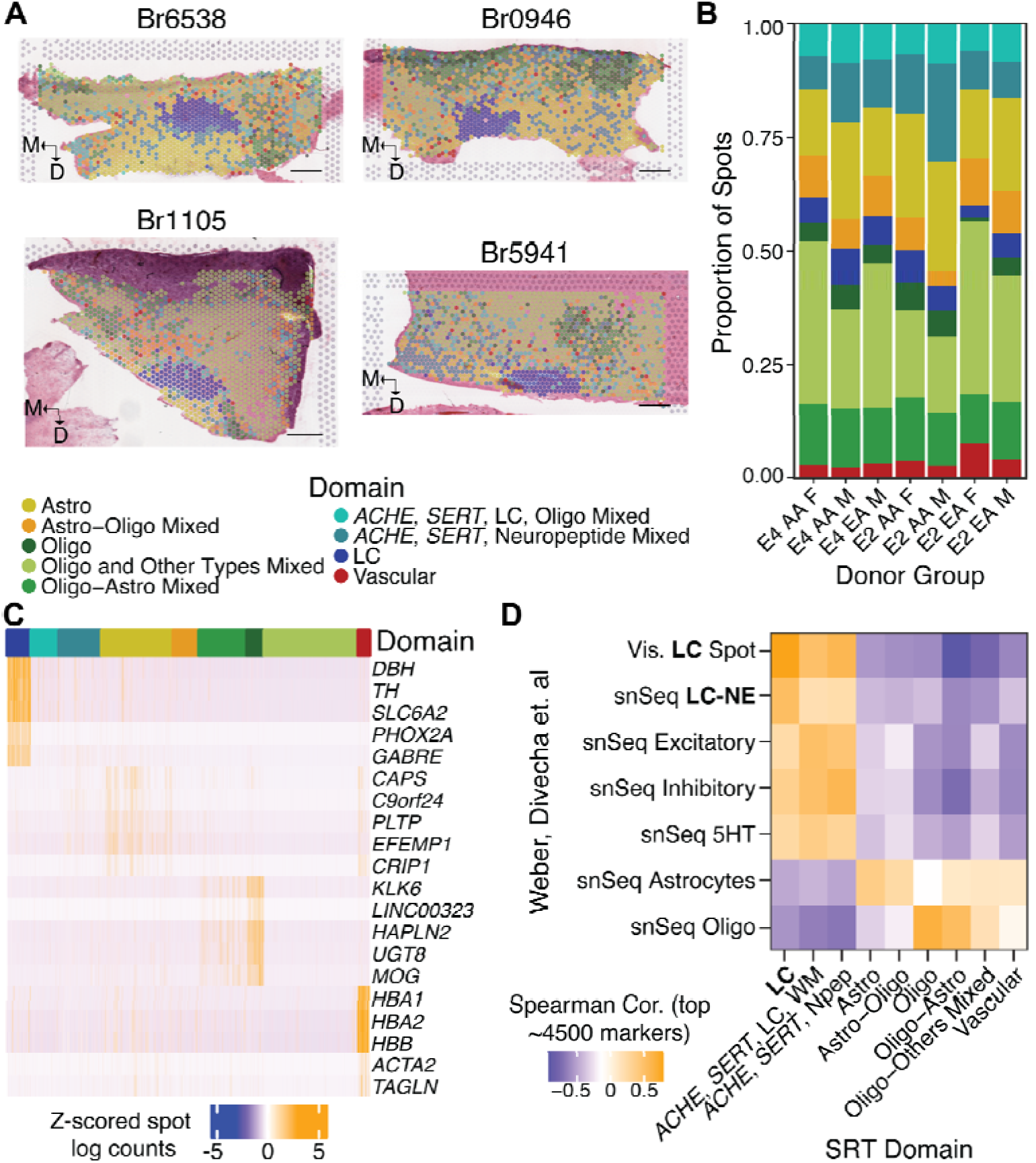
Clustering of spots to generate domains annotated as LC, mixed neurons, and mixed glia. **A)** Tissue sections from 4 representative donors with domain assignments, illustrating that the spatial specificity of the LC d main is consistent across donors and slides. In most instances, the *ACHE*-*SERT-*LC-WM domain is located along one long border of the LC or interspersed within LC; the more scattered *ACHE*-*SERT*-Neuropeptide domain follows a similar pattern in the lower left panel. Arrows indicate the medial-lateral and dorsal-ventral orientations of each tissue section. Scale bars: 500µm. **B)** Proportion of total spots per domain was similar across *APOE* haplotype-ancestry-sex subgroups of donors. Bars are color coded as indicated in panel A. **C)** Top marker genes for each domain are shown on a per-spot basis, with spots grouped by domain color as indicated in panel A. Spot-level visualization illustrates that despite overall enrichment of these genes, they are heterogenously expressed across spots within a domain, suggesting most spots contain multiple cell types. **D)** Registration^88–90^ to supervised annotations of LC/noradrenergic neurons and unsupervised annotations of other cell types from previous SRT and snRNA-seq of the human LC^91^ confirms identity of neuronal and glial spatial domains. Stronger correspondence of the LC domain to prior LC SRT data (as opposed to snRNA-seq) confirms platform-specific identity of the LC domain, likely due to presence of extranuclear transcripts in SRT and cell type-heterogeneity of spots.

Domain proportions were generally similar across donors, *APOE* haplotype, ancestry, and sex (**Fig. 2B**, **Fig. S8**, **Table S3**); however, 4 donors had few or no LC spots despite passing anatomical QC. We determined that subsequent sections collected for SRT were no longer in the LC for 3 of these donors, and they were therefore excluded from downstream analyses (**Fig. S1**, **Table S1**). The LC domain was clearly distinguished by enrichment of canonical marker genes for noradrenergic neurons (*e.g.*, *DBH*, *SLC6A2*; **Fig. 2C**). The remaining domains reflected the heterogenous cell type composition of SRT spots, with five of the 7 other domains enriched in glial markers. This included two domains enriched for canonical astrocyte markers as well as *APOE* (**Fig. 2C**), and three oligodendrocytic domains enriched for two or more genes encoding myelin components (*MBP*, *MOBP*, *MOG*, *MAG*; **Fig. 2C**). We refined the annotations for these glial domains by comparing expression among oligodendrocyte domains or between the astrocyte domains (**Table S4**, *Methods*). Lastly, two domains shared an amalgam of neuronal markers, including the 5-HT reuptake transporter SERT (*SLC6A4*), acetylcholinesterase (*ACHE*), the vesicular glutamate transporter 2 (*SLC17A6*), neuropeptide transmitters including *ADCYAP1*, as well as the 5-HT receptor *HTR2C*. *HTR2C* expression is not documented in transcriptomics from 5-HT neurons^80–83^, suggesting that these domains contain additional cell types. Marker analysis comparing these two domains revealed one domain enriched in LC markers (*DBH*, *SLC6A2*) and myelin genes (*ACHE*-*SERT*-LC-WM; **Fig. 2C**), indicating that this domain likely contains some NE neurons or processes intermingled with neighboring neurons and WM tracts. The other *ACHE*-*SERT* domain was comparatively enriched in several neuropeptides (including *SST*, *TAC1*, and *PENK*) and inhibitory markers *GAD1* and *GAD2* (*ACHE*-*SERT*-Neuropeptides; **Fig. 2C**; **Table S4**). Based on the spatial distributions of these domains, it is possible that they may contain part of the cholinergic laterodorsal tegmental nucleus, which is medial to LC. While peri-LC GABAergic and neuropeptide-only neurons have been described in numerous species^84–86^, neither of these two domains definitively mapped to mouse peri-LC transcriptomic profiles^87^ (**Fig. S9**, **Table S2**).

We performed spatial registration^88–90^ against existing SRT and single nucleus RNA-seq (snRNA-seq) data from neurotypical human LC^91^ to confirm that our annotated domains, particularly the LC domain, were consistent with previous studies. Indeed, our annotated LC spatial domain most strongly corresponded to the LC. Notably, the correspondence to snRNA-seq LC profiles was somewhat lower, likely reflecting both cell type heterogeneity within SRT spots and the unique capability of SRT to recover soma- and neurite-localized transcripts (**Fig. 2D**, **Table S2**).

To better understand the neuronal cell types composing *ACHE*-*SERT* domains, we performed spot deconvolution^92^, including previously defined inhibitory subclusters^93^. A particular subcluster of inhibitory neurons enriched for *CARTPT*, *RGS16*, *CBLN1*, and *SLC18A3* (**Table S5**) appeared to comprise the majority of transcriptomic content in spots from both *ACHE*-*SERT* clusters (**Fig. S10**, **Data S2**). Altogether, our data yielded an unprecedented depth of high-confidence, transcriptome-wide data from LC, along with adjacent neurons, local glia, and WM tracts of the human dorsal pons.

### 2.2 | Transcriptional heterogeneity of the human LC

The human LC is spatially heterogenous in terms of susceptibility to AD pathology, cell structure, and neuropeptide expression^52,53,61–64,94^. Studies in model systems also show spatial heterogeneity across the LC in gene expression, peptidergic cotransmission, axonal projection targets, and cellular physiology^87,95–98^. This heterogeneity is consistent with the ability of the LC to modulate diverse AD-relevant processes including glial activity^9,29–34^, microvascular permeability^26–28^, and cognition^13–22^. It was therefore surprising that we identified a single gene expression-defined LC domain, regardless of how we selected features for dimensionality reduction and performed clustering. To investigate finer molecular heterogeneity, we performed spatially-variable gene (SVG) analysis limited to the LC domain^99^. This analysis identified 27 SVGs at *p*<0.05 in ≥20 samples (**Table S6**). These SVGs included 1) markers of noradrenergic neurons (*TH*, *DBH*, *SLC6A2*), consistent with nonuniform anatomic distribution of these neurons across the LC and adjacent noradrenergic subcoeruleus (*e.g.*^63^), 2) mitochondrial genes, and 3) markers of glia (*MBP*, *GFAP*, *SPARC*) or vasculature (*HBB, HBA2*). Visual inspection revealed that the remaining SVGs were primarily expressed outside of the LC domain with intermediate expression at LC margins. However, *CARTPT* was consistently biased toward mediolateral edges of the LC (medially in some samples and laterally in others; **Fig. S11**). This pattern was not specific to any particular haplotype, sex, or ancestry group. As our annotation of the LC domain was intentionally conservative, it is unlikely that *CARTPT* expression is from other cell types at margins of the LC (**Fig. S11**).

LC heterogeneity also extends to cellular and temporal expression of NM. Up to 40% of large neurons in the adult human LC, including some negative for *TH*, are estimated to be non-pigmented^52^. NM content rises to a plateau in midlife (50-60 years of age) before declining^51^, a process which is accelerated in AD^43^. Therefore, we examined whether gene expression in NM-positive and NM-negative spots of the LC domain may better explain intrinsic and/or AD-associated NM loss. Of the 33 total donors, 29 had robust presence of the LC domain (**Fig. S8**). We used the NM segmentation information in these 29 donors to classify 3,152 NM-positive spots (LC^NM+^; ≥0.3% of spot area containing NM) and 4,763 NM-negative spots (LC^NM-^; **Fig. 3A**). We pseudobulked^100^ LC^NM+^ and LC^NM-^ spots by tissue section and performed differential expression (DE) analysis, controlling for *APOE* haplotype, sex, age, and proportion of AA ancestry (**Table S7**). NM is somatically localized^101^, consistent with the lysosomal structure of NM bodies^56^; it was thus unsurprising that canonical LC marker genes were upregulated in LC^NM+^ spots (**Fig. 3B**), while LC^NM-^ spots were enriched for other cell type markers, including inhibitory neurons (*GAD1* and *GAD2*) and astrocytes (*GJB6*, **Fig. 3B**).

**Figure 3.**
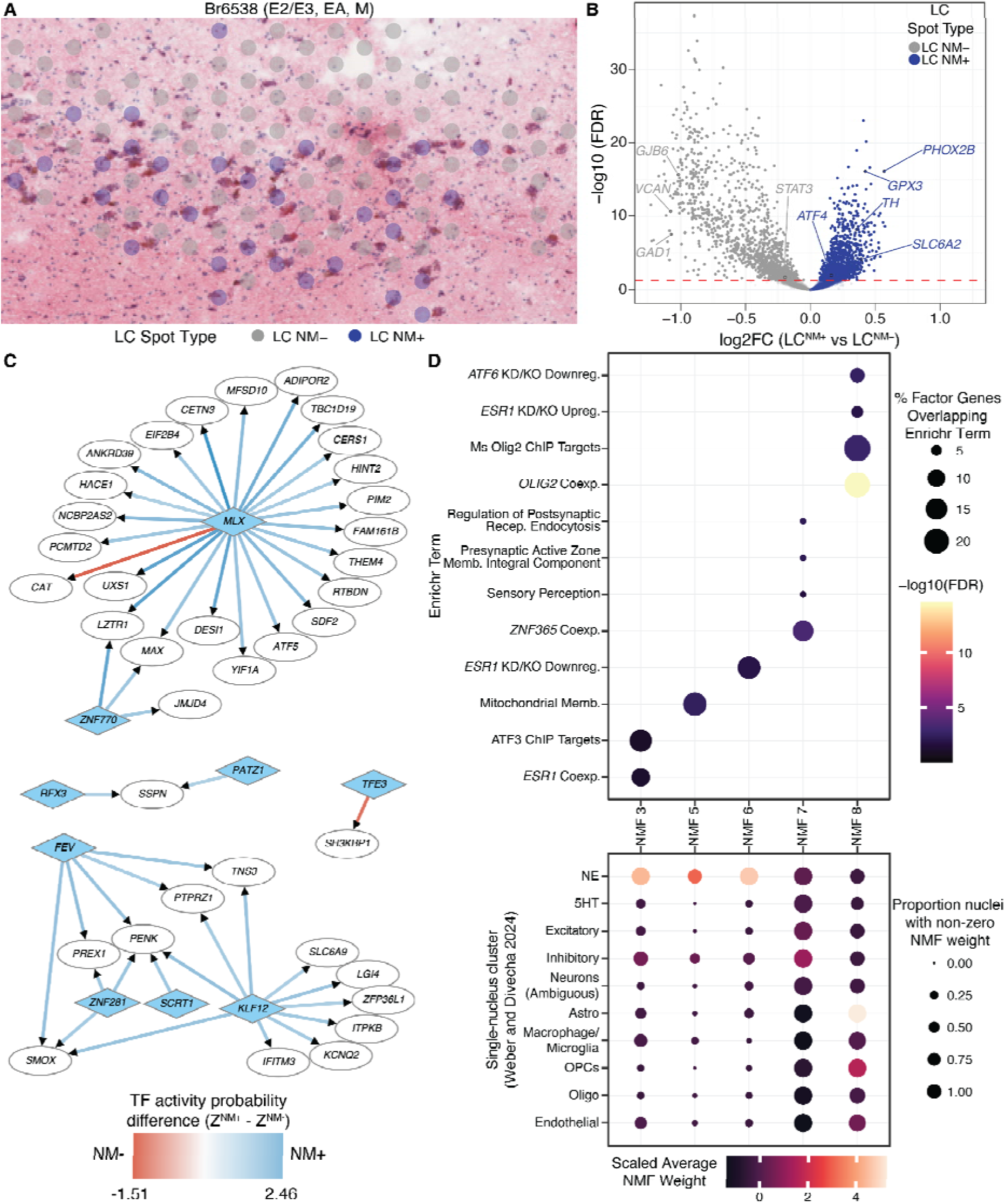
Assessing transcriptional heterogeneity of LC. **A)** NM segmentations of the associated H&E images were used to classify spots in the LC domain as LC^NM+^ or LC^NM-^. Overlaid color labels are size-scaled to SRT spots. **B)** DE analysis between LC^NM+^ and LC^NM-^ spots. **C)** GRN interactions illustrating a subset of TFs (blue diamonds) with high probabilities of regulating target genes (white circles) in LC^NM+^ and LC^NM-^ and differences in those probabilities between the two GRNs (arrow color, legend). **D)** NMF of all LC domain spots resulted in several factors with specific sets of genes. These factor-specific gene sets were tested for enrichment (upper panel) in curated gene sets using *Enrichr*^105^, with the background gene space being the 16.2k genes on which NMF was performed. Projection of the NMF result into our previous snRNA-seq data from human LC^91^ was performed, clarifying the predominant cell type corresponding to each factor (bottom panel).

To better understand the functional implication of these findings, we performed gene set enrichment analyses (GSEA) using MSigDB functional gene sets^102–104^ and sets of transcription factors (TFs) and their putative regulatory target genes from *Enrichr*^105,106^ (**Table S7**). Genes more highly expressed in LC^NM+^ spots were enriched in target genes downregulated by knockout of *TFEB* in human cells (**Table S7**). Conversely, *Tfeb* overexpression in transgenic NM-producing mice significantly increases NM clearance via lysosomal exocytosis^60^. Interestingly, GSEA also indicated LC^NM+^-enriched expression of genes that are regulated by several TFs—*ATF4*, *EBF3*, *STAT3*, and *MEF2D*—all of which are protective against Parkinson’s disease, or associated with NM content in dopamine cells of the substantia nigra^107–111^.

Next, we used two approaches to identify transcriptional programs across the LC domain. First, we used *SCORPION*^112^ to infer separate gene regulatory networks (GRNs) for LC^NM+^ and LC^NM-^ spots. *SCORPION* pseudobulks spots, then uses a message-passing interface (“PANDA”)^113^ to infer the probability of TF-gene regulatory relationships using gene counts and reference datasets of TF-target gene pairs^114^ and TF-TF protein interactions^115^. We compared TF-target (edge) scores between LC^NM+^ and LC^NM-^ GRNs to identify TF-gene interactions likely specific to each subset of LC (**Table S8**, **Data S3**; *Methods*). Many of the largest LC^NM+^-LC^NM-^ edge differences reflected stronger network evidence for LC^NM+^ gene regulation by TFs *KLF12* and *MLX*, a lipid/glucose metabolism-regulated TF^116,117^ (**Fig. 3C**). To complement this, we used a second approach with non-negative matrix factorization (NMF^118^; *Methods*) to identify 160 data-driven NMF factors (**Data S4**). From these, we filtered to factors with uniquely strong weights (≥ 25 genes, *Methods*), which we tested for enrichment in ontology and functional terms (**Table S9**). We separately projected all 160 NMF factors into human LC snRNA-seq data^93^ to identify factors associated with specific cell types (**Fig. 3D**). Interestingly, genes associated with *ATF* family transcription factors were identified in one LC neuron-associated factor (NMF 3) and one astrocyte-associated factor (NMF 6) (**Fig. 3D**). *ATF6* and *ATF4* are broadly involved in neuronal proteostasis responses^119,120^, and *Atf3*- and *Atf6*-regulated genes are induced in response to transgenic expression of NM in the mouse LC^121^. We identified two additional factors associated with LC neurons corresponding to gene regulation by *ESR1* (**Fig. 3D**). Altogether, we demonstrate remarkable molecular heterogeneity in the human LC as a function of anatomical architecture and NM content, including expression of neuropeptides (*CARTPT*), genes regulated by TFs implicated in proteostasis responses and resilience to neurodegeneration, and TF activity.

### 2.3 | Differential expression in human LC across AD risk factors

We next leveraged our LC SRT data to identify gene expression changes that could reflect cellular states protective against or predisposing toward AD development. We therefore pseudobulked^100^ gene expression results by domain to perform differential expression (DE) analyses. As a preprocessing step, we found pseudobulk samples associated with four domains to have too few genes expressed and they were dropped from the DE analysis (**Fig. S12**). Therefore, we included five domains—LC, the two *ACHE*-*SERT* domains, Astro, and Oligo-Astro—from *n*=29-30 donors (**Table S1**) in the DE analysis, controlling for age, sex, *APOE* haplotype, and genomic ancestry proportion.

#### 2.3.1 | Sex-specific differential expression in human LC associated with AD

Because females are more frequently diagnosed with and more severely affected by AD^65–67^, we began by examining the transcriptomic effects of sex on the LC and surrounding tissue. Remarkably, all five domains showed robust sex differences in gene expression, ranging from 523 to 1,493 sex-DE genes (DEGs, FDR < 0.05, **Fig. S13**, **Table S10**). In the LC domain, male upregulation was observed for genes including 1) *CHGB*, a core component of adrenal catecholamine vesicles whose loss impairs uptake of catecholamine precursor L-DOPA and reduces vesicular epinephrine—but not NE—content/release^122^, and 2) *PRCP* (prolylcarboxypeptidase), an activator/inactivator of peptide hormones, such as the *POMC*-derived metabolic hormone α-MSH^123^ (**Fig. 4A**). Notably, *POMC* was itself only expressed in the LC and ACHE-SERT-LC-Oligo domains. In females, upregulation of *APOLD1*—a regulator of endothelial permeability^124^ —and glutathione peroxidase *GSTT2B* was observed in the LC domain. Additionally, the transcriptional cofactor *TCF20* was significantly female-upregulated, as were regulatory targets of TCF20 (**Fig. 4A**, **Table S11**). Female-upregulated sex-DEGs in the Astro domain were notable for roles in autophagy, the golgi network, and protein folding (*HSPB8*, *GASK1B*, *DNAJB1*, respectively; **Fig. 4B**). Interestingly, in the *ACHE*-*SERT*-LC-Oligo domain, the autophagy-regulating partner of *HSPB8*, *BAG3*, was male-upregulated, along with *PHYH* (peroxisomal degradation of 3-methyl fatty acids; **Fig. 4C**). In females, the *ACHE*-*SERT*-LC-Oligo domain showed greater expression of *CPNE6*, which inhibits spontaneous neurotransmitter vesicle release at presynaptic sites in a calcium-dependent manner^125^ (**Fig. 4C**). Altogether, these findings suggest that sex strongly impacts expression of genes critical to degradative and exocytotic cellular compartments (phagosomes, peroxisomes, neurotransmitter vesicles) and protein folding across LC, astrocytes, and other mixed neuronal cell types.

**Figure 4.**
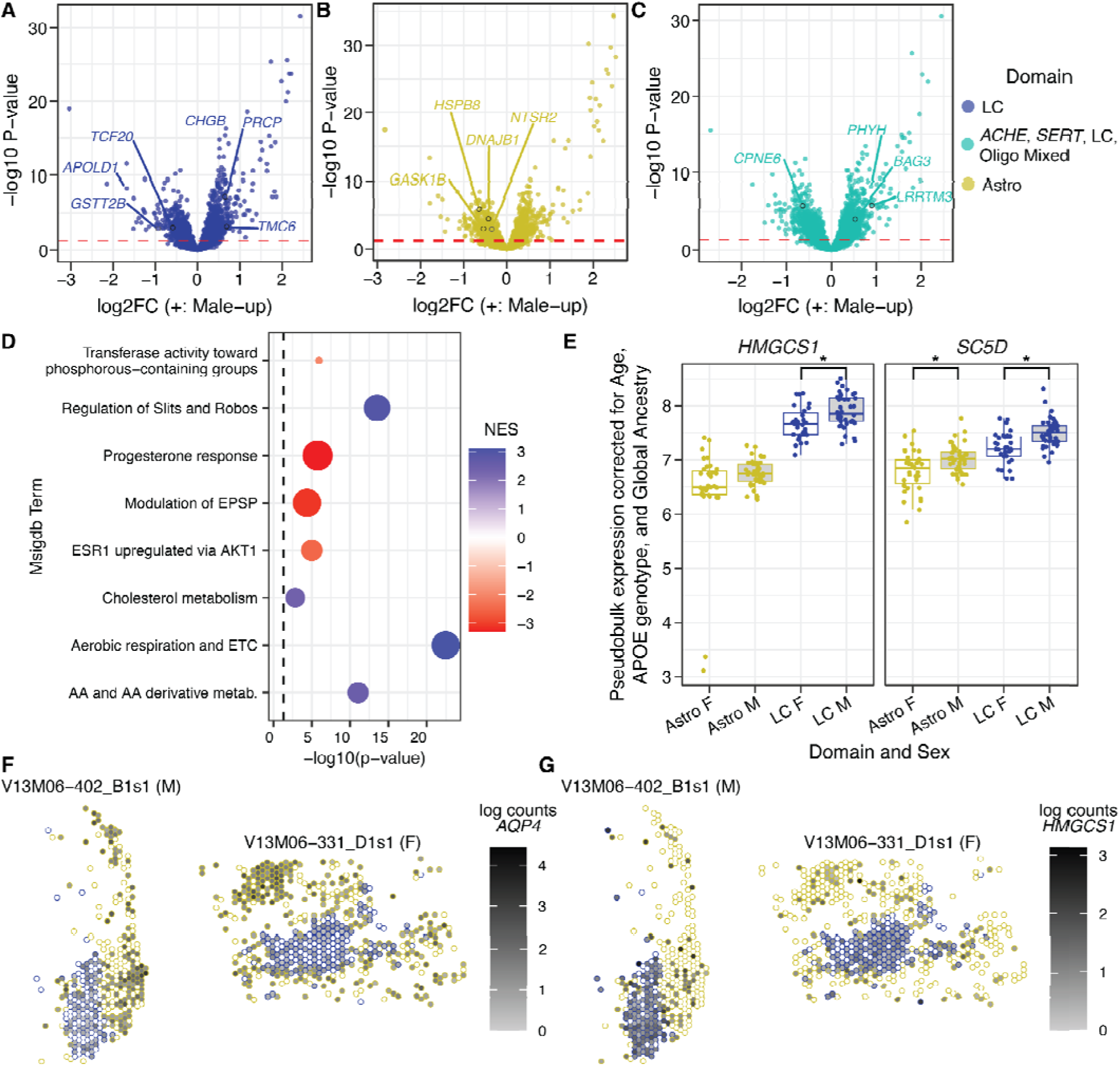
Sex DE in LC, surrounding neurons, and astrocytes. **A)** Volcano plot illustrating sex DE in the LC domain. Red line indicates an FDR threshold of 0.05. **B)** Volcano plot illustrating sex DE in the Astro domain. Red line indicates an FDR threshold of 0.05. **C)** Volcano plot illustrating sex DE in the *ACHE*, *SERT*, LC, and WM mixed domain. Red line indicates an FDR threshold of 0.05. **D)** GSEA results for sex-DEGs in the LC, with the sign of the enrichment score (NES, normalized enrichment score) indicating enrichment among male-upregulated (blue) or female-upregulated (red) genes. GSEA results for all domains are available in **Table S11**. **E)** Pseudobulk expression per sample of selected cholesterol synthesis pathway genes in the LC domain and Astro domain, stratified by sex. * denotes FDR<0.05. **F-G)** As cholesterol synthesis genes are generally astrocyte-expressed, we visualized expression in the Astro (yellow) and LC (blue) domains to compare expression patterns of a canonical astrocyte-specific gene, *AQP4* (**F**), alongside cholesterol synthesis gene *HMGCS1* (**G**) in 2 representative donors (Br1884 and Br5426).

To understand the functional implications of sex-DE, we used GSEA^102,126^ to test for enrichment of male- and female-biased DE in gene sets spanning ontologies and pathways defined by MSigDB^102–104^. We similarly tested for enrichment of DE in target genes of TFs using collected TF-target sets^105,106^ as previously described^127^ (**Table S11**). Focusing on sex-DE in the LC domain, there was a clear enrichment of sex hormone-regulated genes among female-biased genes, while male LC was enriched for terms relating to metabolic and mitochondrial processes (**Fig. 4D**). Though the enrichment was modest in terms of significance, we were struck by male-enriched expression of cholesterol metabolism genes (**Fig. 4D**), which included sex-DE of several cholesterol synthesis pathway genes such as *HMGCS1* and *SC5D* (**Fig. 4E**). Cholesterol synthesis is generally localized to astrocytes, and visualization of *AQP4,* a canonical astrocyte marker, revealed modest expression in most LC domain spots, but strong expression in Astro domain spots (**Fig. 4F**). By contrast, cholesterol synthesis genes were expressed more highly and densely in the LC domain relative to the Astro domain in both sexes (**Fig. 4E,G**). These findings suggest that the LC may have capacity to synthesize cholesterol locally, and that this process may be more active in the LC of males.

#### 2.3.2 | *APOE* allele-specific differential expression in human LC associated with AD

The largest-effect common genetic risk factor for AD is in the *APOE* locus, where the most common variant, E3, is neutral, while E4 substantially increases AD risk and E2 is protective^69,72^. Importantly, a smaller degree of risk is conferred by E4 in African ancestry (AA) individuals relative to European ancestry (EA) individuals^68,69^. As our cohort was balanced for carriers of E4 or E2 (both heterozygotes and homozygotes) across these two genomic ancestry groups, we first examined whether *APOE* haplotype alters the transcriptome (haplotype DE) of the middle-aged LC and surrounding domains. Additional (ancestry-controlled) DE analyses between diplotypes in our dataset (E2/E2, E2/E3, E3/E4, E4/E4) and certain diplotype combinations (*e.g.*, E4 homozygotes vs. all E2 carriers) were also performed (**Table S10**) given the profound effect of E4 homozygosity on AD^72^. We then extended these analyses to assess whether E4-E2 differences are more widespread in EA individuals (ancestry-haplotype DE). Given the low frequency of E2 and E4 alleles, we prioritized the best-powered analyses within and across ancestry groups.

We assessed haplotype DE (*n*=29-30) in the five domains described above while controlling for age, sex, and AA genomic ancestry proportion as a continuous measure (**Fig. S14**, **Table S10**). In the LC domain, E4 carriers showed comparatively lower expression of canonical astrocyte genes, such as *SLC1A3* and *GJA1* (**Fig. 5A**), as well as the neurotensin receptor *NTSR2*. In the Astro domain significant downregulation of heat-shock proteins—essential to preventing protein misfolding—was observed in E4 carriers (**Fig. 5A**). We then performed GSEA^102,126^ for MsigDB^102–104^ and TF-target gene sets (**Table S11**), which highlighted downregulation of multiple sets of astrocyte markers in the LC domain of E4 carriers (**Fig. 5B**). By contrast, cell cycle terms—which would not be expected in healthy neurons—were enriched for E4-upregulated genes. Previous work shows genes associated with aberrant cell cycle re-entry in other human brain regions^128^ become dysregulated in human neurons, including LC neurons, during aging and AD. Alternatively, these cell cycle terms may be attributable to cells adjacent to LC neurons with bonafide mitotic potential (e.g., microglia). For example, cells dividing in the LC domain of E4 carriers could displace LC-localized astrocytes found in E2 carriers, resulting in a net decrease in astrocyte gene expression in LC. Altogether, these findings suggest that *APOE* haplotype-specific differences emerge in close proximity to LC neurons, such that local astrocytic cell density or expression differs between E2 and E4 haplotypes.

**Figure 5.**
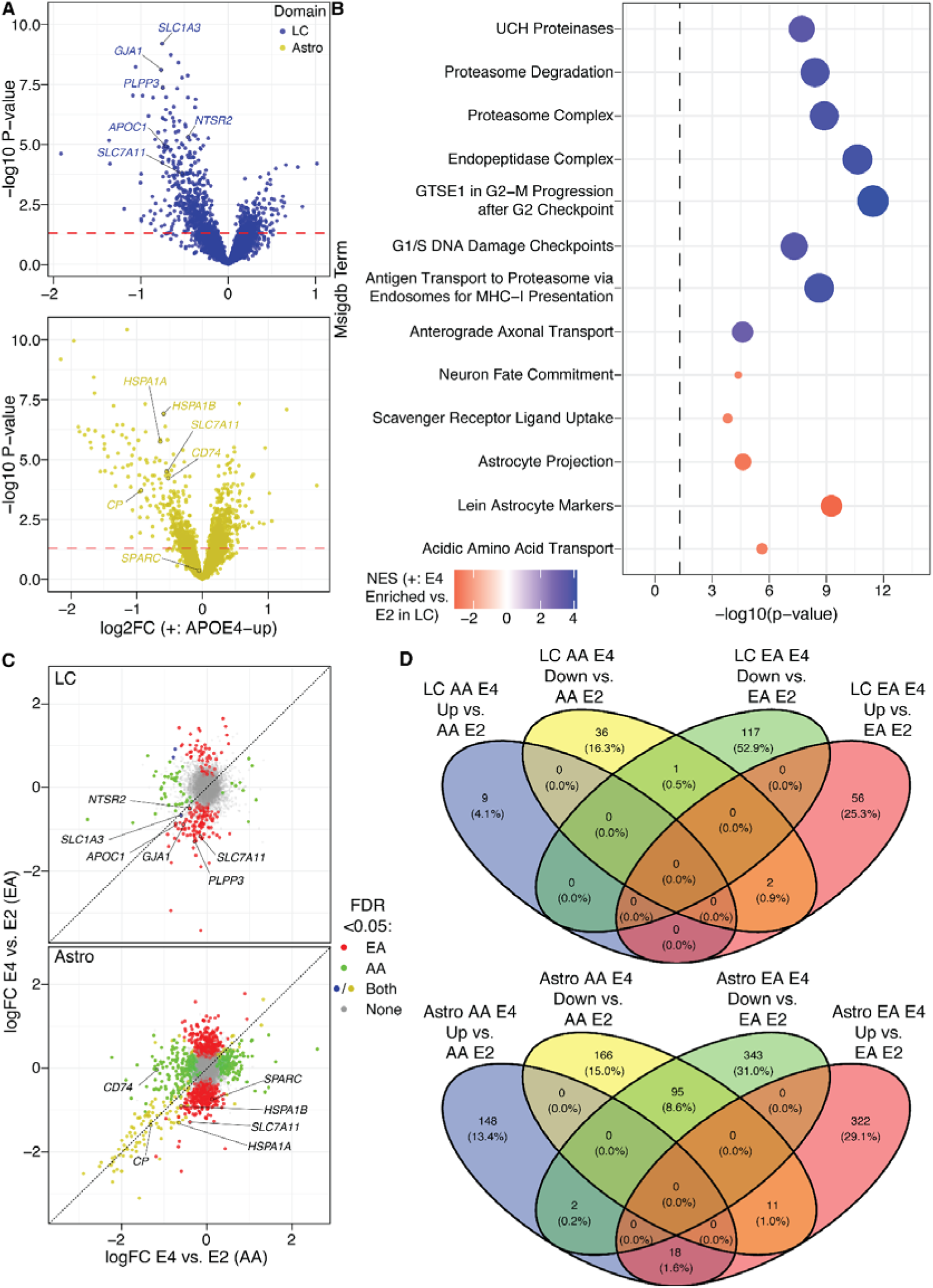
Haplotype DE in LC and surrounding astrocytes between, collectively or stratified by predominant genomic ancestry. **A)** *APOE* haplotype DE results between E4 and E2 carriers in the LC and Astro domains (N=29 and 30, respectively). **B)** GSEA terms enriched in genes with greater LC expression in E4 or E2 carriers. **C)** Comparison of log fold-change (logFC) in LC and Astro domains from ancestry-haplotype DE analyses. Points are colored to indicat DEGs identified in one or both ancestries. **D)** Venn diagrams illustrating that the vast majority of haplotype DE in LC and Astro is ancestry-specific, with more haplotype DEGs in EA.

#### 2.3.3 | Genetic ancestry-specific differential expression in human LC associated with AD

The clinical impacts of *APOE* haplotype vary by ancestry, with E4 having a smaller risk effect in AA individuals relative to EA, while the protective effect of E2 is similar across these ancestry groups^69^. We thus asked whether LC gene expression was affected by *APOE* haplotype in an ancestry-differentiated manner—and whether EA E4-E2 comparisons showed broader effects relative to AA E4-E2 comparisons, which would support an etiologically important role of *APOE* haplotype in LC vulnerability to AD pathology. We therefore stratified donors into groups of ≥75% genomic EA (*n*=10) or AA (*n*=16) to analyze ancestry-haplotype DE (**Fig. S15**-**Fig. S18**, **Table S1**; **Table S10**) and identified MSigDB^102–104^ GSEA^102,126^ enrichments unique to haplotype-DE in one ancestry group (**Table S12**). In the LC, AA haplotype-DEGs (E4-upregulated) were uniquely enriched for 26S proteasome-mediated protein degradation, while E4-upregulated EA haplotype-DEGs were uniquely enriched for mitochondrial and oxidative phosphorylation terms. In the Astro domain, E4-upregulated genes were enriched in EA for kinesins, microtubule-based transport, and retrograde golgi-to-endoplasmic reticulum transport; synaptic and dendritic terms were enriched among E4-upregulated genes in both ancestry groups. E2-upregulated (E4-downregulated) DEGs were enriched in differing sets of putative regulatory targets of *ATF6*, a proteostasis TF (**Table S12**), between the ancestry groups.

However, the ancestry-haplotype DE results were perhaps most striking in their exclusivity: in the LC domain, ancestry-haplotype DEGs were mostly unique to EA (**Fig. 5C-D, upper panels**). A similar pattern was observed in the Astro domain, with most E4-upregulated genes relative to E2 being distinct between ancestries (**Fig. 5C-D, lower panels**). Nonetheless, a small set of ancestry-haplotype DEGs were common to EA and AA: in the LC domain, astrocytic genes such as *SLC1A3* and *GJA1* showed similar log fold-changes (logFCs) in both ancestries (despite only being significant in one ancestry group each), suggesting a common relationship between astrocytes, LC neurons, and *APOE* haplotype. In the Astro domain, a shared group of genes such as ceruloplasmin (*CP*) was significantly downregulated at relatively large magnitudes (|logFC| > 1) in E4 relative to E2 carriers; this ancestry-shared set of E4-downregulated Astro DEGs also included hemoglobins and 7 ciliary and flagella associated protein (*CFAP*) family genes, potentially indicating an overall decrease in astrocyte processes and/or astrocyte-vascular interfaces in E4 carriers of both ancestries. Altogether, These findings suggest that while gene-level effects of *APOE* haplotype on LC and dorsal pontine astrocyte gene expression are largely unshared between ancestries, 1) E4 consistently downregulates a set of astrocytic genes relative to E2, 2) E2 carriers of both ancestries display a common relationship between the LC and embedded astrocytes, and 3) *ATF6*-regulated genes are differentially expressed between E4 and E2 carriers in pontine astrocytes. Moreover, clinically observed differences in AD risk conferred by the interaction of E4 carrier status and ancestry are recapitulated at the molecular level in our DE observations in LC. Particularly, we observe a greater breadth of E4-E2 differential expression in EA relative to AA individuals, consistent with an etiologic role of LC in *APOE*-mediated AD risk.

### 2.4 | Gene expression associations with LC NM implicate *APOE* and neurodegeneration-relevant pathways

NM-sensitive MRI has demonstrated that LC depigmentation has predictive value for rate of cognitive decline, symptom severity, and pathology burden in AD^40,43,129^. NM content varies from neuron to neuron^39^ and is dogmatically considered to be permanent until cell death^56^, though some studies suggest intrinsic clearance may occur at a slow rate in intact neurons^39,59,60^. Moreover, LC neurons with the highest NM content are preferentially lost in PD ^130^, suggesting that dysregulated NM homeostasis plays a direct role in susceptibility of LC neurons to neurodegeneration. While our above investigation of transcriptomic heterogeneity across the entire LC domain contrasted LC^NM+^ and LC^NM-^ microenvironments, we also sought to investigate whether NM pigmentation within LC^NM+^ spots corresponds to transcriptional states. We therefore linear mixed modelling between spot-level gene expression and NM intensity, where darker pixels signify more pigment. While only LC^NM+^ pixels were used to calculate NM intensity, it should be noted that associations with gene expression may reflect other cell types within the spot modulating NM intensity—for example, via glia-LC signaling or inflammatory responses of microglia toward extracellular NM^57,131^. An additional analysis using LC^NM+^ pixel proportion, the percentage of each LC SRT spot covered by NM, was also performed (**Table S13**). This analysis showed that spots covered by greater areas of NM had greater expression of LC marker genes, and those with less NM had greater expression of other cell type markers, consistent with NM-positive spots being enriched for NE neurons.

Modeling of NM intensity associations with expression of 23,744 genes yielded 809 significant genes (FDR < 0.05) when controlling for E4/E2 carrier status, sex, age, and genomic ancestry. We observed a surprising inverse association of *APOE* gene expression with NM intensity, with greater *APOE* expression associated with less pigmented, or lighter NM (**Fig. 6A**). Expression of several antioxidant metallothioneins (*MT2A*, *MT1E*, *MT1X*, *MT3*) also showed significant inverse relationships with NM intensity (**Table S13**), which is consistent with NM production as an oxidative process and with NM bodies containing metal ions such as iron and copper. Meanwhile, darker NM was associated with greater expression of genes involved in protection against oxidative stress, including *GPX3* (FDR<7.0x10^-12^) (**Fig. 6B**). Several additional associations between higher gene expression and darker NM were noteworthy, including *SARAF* (Bonferroni FDR<1.3x10^-4^; **Fig. S19**), which encodes a protein that halts cytosolic-to-endoplasmic reticulum (ER) Ca^2+^ repletion once ER stores are adequate^132^. We also found increased expression of the autophagosome biogenesis gene *MAP1LC3B* (FDR<2.5x10^-7^, **Fig. S19**) in darker NM spots. Notably, autophagosomes deliver their cargo to lysosomes, double-membraned organelles closely resembling NM granules^56^. Finally, darker NM was associated with higher expression of *PGRMC1* (FDR<4.5x10^-10^, **Fig. S19**), which encodes a protein acting as both a heme chaperone and facilitator of LDL receptor (LDLR) activity^133,134^.

**Figure 6.**
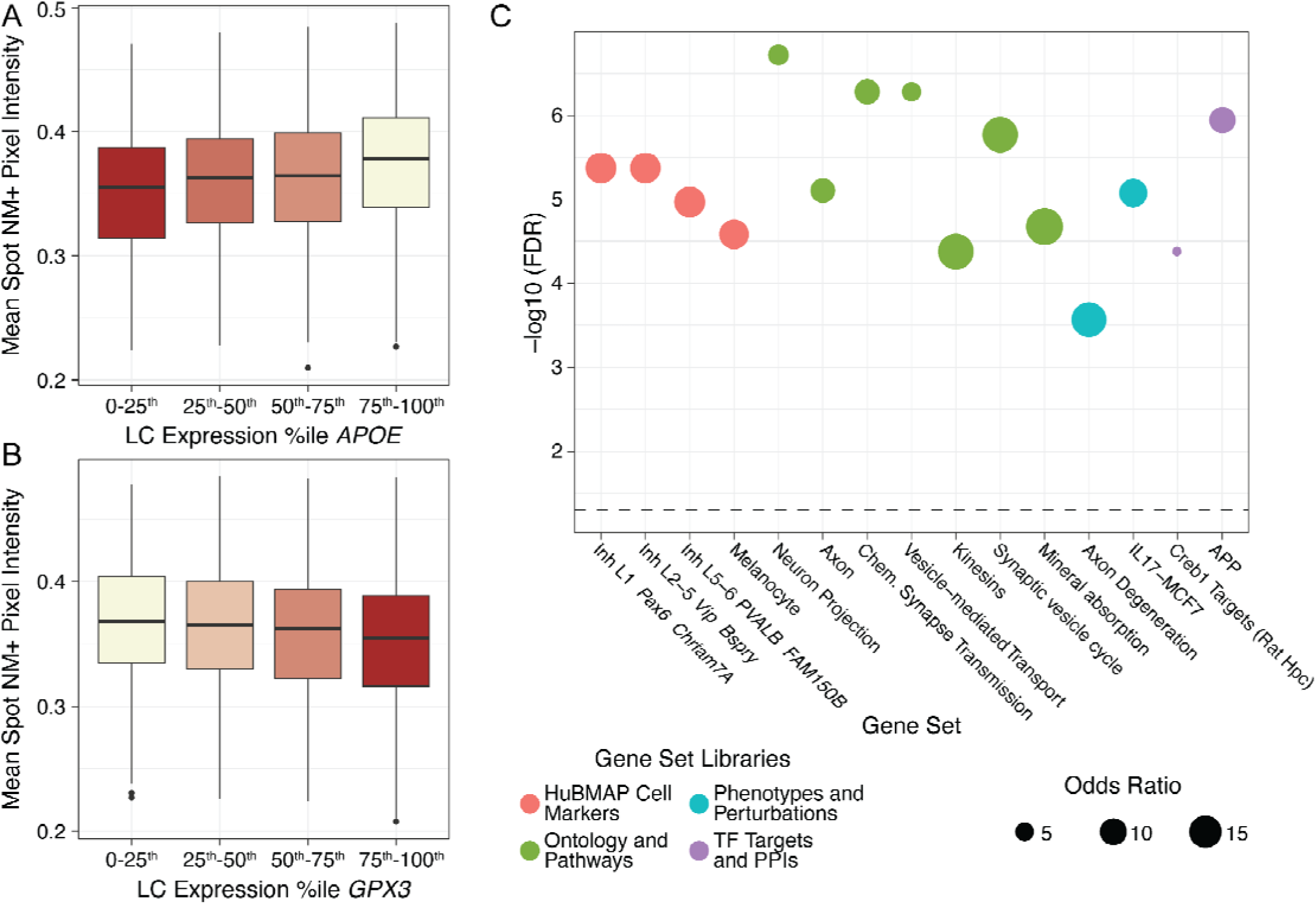
NM intensity is associated with gene expression in the human LC. **A-B)** Spot means of NM pixel intensities within LC^NM+^ spots are plotted against spot-level nonzero expression quartiles of *APOE* or *GPX3* from all samples containing ≥20 LC^NM+^ spots. Note that NM intensity values range 0 to 1, with 0 being darker/most pigmented. **C)** *Enrichr*^105^ analysis of all genes associated (Bonferroni *p*<0.05) with LC^NM^.

To better understand the biological role of LC^NM^-associated genes, we used *Enrichr*^105^ to identify potential pathways, ontologies, and gene regulatory signatures that may influence NM content (**Table S14**, **Data S5**). Melanocyte gene markers were enriched among LC^NM^-associated genes, suggesting common biology of peripheral and central pigment processing (**Fig. 6C**). Interestingly, LC^NM^-associated genes w re enriched for inhibitory neuron-specific ontology terms, suggesting potential regulation of NM production by local inhibitory signaling. There were also strong enrichments of LC^NM^-associated genes in neurodegeneration-relevant pathways, including protein interactors of amyloid-ß precursor protein (APP) and degeneration of axons, which are the first subcellular structure to accumulate pTau in human LC^1^. Altogether, these findings highlight that NM content in the LC is associated with AD risk factors (*APOE*), lipid production and transport, calcium flux, oxidative stress, AD-associated biological complexes, and early degenerative processes.

## 3 | Discussion

The LC is one of the earliest sites of AD-associated pTau deposition—occurring near-universally in adulthood and exacerbated in AD^1^—resulting in substantial accumulation well before clinical symptoms manifest. In aging and the early stages of AD, the LC shows hyperexcitability based on NE and NE metabolite levels^6,7,136^, potentially underlying symptoms of early/prodromal AD^2,41,137^. Simultaneously, early AD is associated with a decline in NM pigmentation of the LC^39,43^, with neuroimaging studies further demonstrating that LC NM content is associated with AD symptoms^41^ and prognosis^43^. Similarly, LC pTau burden predicts subsequent pTau accumulation in interconnected vulnerable brain areas including the entorhinal cortex^40^. The LC thus constitutes an ideal target for early interventions to treat AD symptoms and potentially mitigate the spread of AD pathology across the brain. To better understand the impact of AD-associated risk on LC biology prior to pTau’s ascending spread and overt LC degeneration, we performed SRT from 33 neurotypical donors without pathological AD balanced for epidemiologic AD risk factors including sex, *APOE* haplotypes, and genomic ancestry. We stringently defined the LC domain using the resulting gene expression and histological images, and leveraged the data to quantify the molecular impacts of AD-associated risk factors on the LC and local pons.

While the rodent LC shows striking spatial heterogeneity across molecular, cellular, physiological, and circuit properties, a paucity of studies in the human brain has left a significant gap in our understanding of the molecular and cellular organization of the human LC. We therefore leveraged our data to directly investigate heterogeneity at the molecular and cellular levels in the human LC. Previous reports have estimated that up to 40% of LC neurons are NM-negative^52^, which is important to consider in the context of NM loss via LC degeneration in aging^51^ and AD. We performed DE analyses to determine whether LC^NM+^ and LC^NM-^ subdomains constituted distinct niches of LC neurons. LC^NM+^ spots were mildly enriched for canonical LC marker genes such as *DBH* and *TH* relative to LC^NM-^. Although NM loss^39^ precedes death of LC neurons, LC^NM-^ spots were enriched for markers of altogether different cell types. We interpret this to reflect LC^NM-^ spots containing more space for NM-negative neuronal processes to intermix with other cell types compared to LC^NM+^ spots, as NM localizes to the large somata of NE neurons which presumably occupy more space within each spot^88,91^. Nonetheless, GSEA^102,126^ revealed that LC^NM+^-enriched genes were overrepresented in targets of multiple TFs (*ATF4*, *EBF3*, *STAT3*, *MEF2D*; **Table S7**) previously associated with NM and PD resilience in the substantia nigra^107–111^. This suggests that NM induces or is coexpressed with pro-resilience factors in neurodegenerative disease. We next compared gene-regulatory networks (GRNs) of LC^NM+^ and LC^NM-^ and found that the TF *MLX,* which is associated with hepatic *de novo* lipid synthesis^116,138^, was more likely to regulate several genes in LC^NM+^ including *ADIPOR2*, a receptor for the metabolic hormone adiponectin. Finally, we applied NMF^118^ to identify discrete gene coexpression patterns (factors) of LC neurons and other cell types contained within LC domain spots. Two NMF factors—one driven by LC neurons and one by astrocytes—were uniquely defined by sets of genes implicating estrogen-regulated expression. These findings suggest roles for sex in molecular heterogeneity of the human LC, consistent with previously observed findings in rodents^80,87^. While studies at cellular resolution will be needed to more clearly delineate spatial and molecular niches, our findings provide evidence for heterogeneity at the molecular level in the human LC.

Given our findings showing coexpression of estrogen-regulated genes LC and Astro domains, we examined transcriptomic differences between sexes in the LC and surrounding domains (females, *n*=31-35 samples, *i.e.*, tissue sections, per domain from 10-11 donors; males, *n*=42 samples per domain from 19 donors). As LC excitability increases not only in early AD, but also in healthy aging^139,140^, it is interesting that female-biased LC sex-DEGs showed enrichment in genes associated with “modulation of [excitatory postsynaptic potentials]”, suggesting a possible sex difference in excitability of the aging LC. Interestingly, male-biased expression was enriched for oxidative phosphorylation and cholesterol metabolism, general metabolic processes downstream of *MLX* that were implicated in our LC^NM+^ gene regulatory network. Astrocytes are established as the primary site of cholesterol synthesis in the brain^141,142^, but we found that genes specifically involved in *de novo* cholesterol synthesis were highly expressed in an LC-localized manner. This surprising feature of LC biology cannot be explained by its extensive cell membranes and cholesterol-rich lipid rafts, as the human LC is enriched in cholesterol synthesis genes relative to the substantia nigra, another catecholaminergic projection nucleus^143^. These data raise the interesting possibility that the capacity of LC neurons to synthesize cholesterol, rather than obtain it via uptake of *APOE*-coated lipoproteins, reduces AD risk. Our data further invite the question of whether such an effect contributes to the lower AD risk in males.

To investigate the impact of *APOE* haplotype on the dorsal pontine transcriptome, we performed DE analysis within each domain between E4 and E2 carriers (E4, *n*=49 samples from 19 donors; E2, *n*=24-28 samples from 10-11 donors). Nonetheless, both astrocytes and LC showed substantially different expression profiles across *APOE* haplotypes. Strikingly, some of the largest E4 effects in the LC domain included downregulation of canonical astrocyte genes such as *SLC1A3* and *GJA1*. The limitations of spot-based SRT should be considered when evaluating this finding, which we interpret to signify APOE-mediated effects on the abundance or transcriptional state of astrocytes neighboring LC neurons. However, cellular-resolution studies will be required to better understand the changes in LC-astrocyte interactions in E4 carriers.

While E4 is an AD risk factor—and E2 a protective factor—in both AA and EA individuals, only the effect of E4 substantially differs between these ancestry groups^69^. Therefore, if the LC is etiologically important to AD, the molecular effects of *APOE* haplotype should recapitulate the observed ancestry differences in E4-phenotypic outcome associations. Indeed, we observed more E4-E2 DEGs in the LC of EA carriers than in AA when we stratified our cohort by genomic ancestry (EA, *n*=20-24 samples from 9-10 donors; AA, *n*=41 samples from 16 donors). Moreover, nearly all gene dysregulation by E4 was ancestry-specific in the LC (**Fig. 5C-D**, upper panels). While the Astro domain also showed ample ancestry-specific *APOE* effects, genes markedly downregulated in E4 donors were common to both ancestries (**Fig. 5C-D**, lower panels), suggesting that E4-downregulatory effects on pontine astrocytes may be ancestry-nonspecific.

NM-sensitive MRI of the LC shows significant promise as a biomarker for AD prognosis and progression^41,43,44^; however, little is known about how NM impacts LC function in aging or AD. We leveraged the ability of SRT to provide quantitative histological data and transcriptomic data from the same sample to assess associations between NM content and LC gene expression at the spot-level. NM-associated genes were relevant to several aspects of LC biology and AD-associated LC dysregulation. First, lighter NM pigmentation was associated with several antioxidant metallothioneins (*MT2A*, *MT1E*, *MT1X*, *MT3*), consistent with catecholamine oxidation by—and NM enrichment in—(free) iron^49^. This finding suggests that iron/copper sequestration by metallothioneins may disrupt oxidative production of NM from NE. However, genes associated with lighter NM could also reflect NM clearance processes: NM appears to decrease in dopamine and NE neurons of AD subjects^39,59^, and NM-producing mouse experiments^60^ may suggest the existence of a regulated NM clearance mechanism. Meanwhile, darker NM was associated with glutathione peroxidase *GPX3* expression, consistent with darker NM content associated with the related gene *GPX4* in the substantia nigra^144^.

Additional domain marker genes and NM-associated genes could potentially link NM with aging/AD-associated LC hyperexcitability and elevated NE release. Marker analysis across the three WM-enriched domains revealed the Oligo domain to be enriched for amyloid precursor-related peptide *APLP1*, which may promote internalization of the noradrenergic autoreceptor α2A (*ADRA2A*)^145^. This suggests a possible interaction between LC neurons and subpopulations of local oligodendrocytes that impair feedback inhibition of NE release. Meanwhile, darker NM was associated with greater expression of *SARAF*, whose protein product halts cytosolic-to-endoplasmic reticulum (ER) Ca^2+^ transfer once ER stores are replenished^132^. Dysregulation of intracellular Ca^2+^ can alter neuronal excitability or trigger apoptosis. Darker NM was also associated with greater expression of *MAP1LC3B*, a marker of autophagocytosed neurotransmitter vesicles, which in LC neurons would contain DBH and its copper cofactor along with NE (*i.e.*, NM precursors). Impaired vesicle autophagocytosis in rodent hippocampal axons increases neuronal excitability^146^. Though cytosolic (rather than vesicular) catecholamines form NM in mouse substantia nigra^147^, this has not been demonstrated in the LC or in human cells; further, lysosomes (*i.e.*, mature NM bodies^56^) receiving both cytosolic NE and noradrenergic vesicles would ultimately contain the major components of NM. Darker NM was also associated with higher expression of genes implicated in lipid/sterol biology. *PGRMC1* encodes a protein acting as a heme chaperone and facilitator of LDL receptor (LDLR) activity. Given our findings regarding cholesterol synthesis and activity of a *de novo* lipogenesis regulator (*MLX*) in LC—along with the NM’s enrichment in iron—this finding further implies a relationship between lipids/sterols and NM or its iron content. As oxidative stress is propagated within cells via lipid peroxidation, these genes suggest NM content may index (buffering capacity against) oxidative stress. Finally, NM intensity-associated genes were enriched for inhibitory neuron markers. Though we were unable to define a specific domain comprised only of peri-LC inhibitory neurons^84–87^, enrichment of canonical GABAergic markers (*GAD1* and *GAD2*) alongside inhibitory neuropeptide co-transmitters (*e.g.*, *ADCYAP1* and/or *SST*) suggested these cells are present in the *ACHE*-*SERT* domains. Because these neurons provide direct inhibitory input to the LC, they carry significant potential to modulate LC function and excitability in neurodegenerative disease. Future high resolution studies in postmortem human LC should emphasize identifying these neurons and understanding how AD risk factors impact their transcription.

Our analyses broadly identify two recurring themes: lipid/sterol pathways and astrocytic gene expression in LC microenvironments as a trans-ancestry effect of *APOE*. First, the LC shows unexpected expression of lipid/sterol synthesis genes alongside inferred NM-specific transcriptional regulation via a metabolically-controlled TF, *MLX*, targeting genes including metabolic signal receptor *ADIPOR2*. AD subjects show reduced cholesterol and downregulation of cholesterol synthesis genes in the entorhinal cortex and hippocampus, brain regions that are innervated by LC and have high pTau burdens in early AD^148^. However, neuronal cholesterol may also facilitate formation of PD-associated Lewy bodies from alpha-synuclein^149^. Our finding that cholesterol synthesis gene expression is male-biased in the LC provides a potential mechanism of AD resilience and PD susceptibility that could explain their opposing sex differences in incidence^150^. It is additionally noteworthy that NM is rich in specific lipid families^56^; our findings may indicate these lipid synthesis pathways are activated to replenish membranes used to envelop NM. The association between LC degeneration/NM dissipation with AD risk and progression supports a model where sustained NM production is protective against AD, and we thus surmise that maintenance of LC neuronal lipid/sterol self-sufficiency would also be protective.

Second, a consistent and unexpected set of findings highlighted *APOE* haplotypes impacting expression of astrocytic genes within the LC domain. As astrocytes in the LC domain are necessarily close to LC somata or processes, this suggests that astrocyte-LC interactions are perturbed by *APOE* haplotype. Specifically, the increased astrocytic marker expression within LC of E2 carriers suggests that astrocyte density or transcriptional state proximal to LC neurons contributes to the protective effect of the E2 haplotype. However, further studies at cellular resolution are necessary to understand these interactions, their differences in E2/E4 carriers, and how they mechanistically contribute to, or protect against, AD. Importantly, the enrichment of canonical astrocyte markers among E2-upregulated (*i.e.*, E4-downregulated) genes in the LC domain was consistent across ancestry (**Table S10**). As E4, but not E2, shows differing effects on clinical AD in EA/AA^69^, we infer that our observations regarding *APOE* haplotype effects on astrocytic genes in the LC are likely an E2-mediated, AD-protective phenomenon. Thus, modulating the localization or state of LC-proximal astrocytes could be a therapeutic strategy whose efficacy would be consistent across EA and AA individuals. While the astrocytic component of the LC microenvironment (and downregulated genes in the Astro domain itself) appear similarly affected by *APOE* haplotype across AA and EA, our observations also indicate that other cell types are affected by E4/E2 in a highly ancestry-specific manner. These ancestry-specific findings provide further insights into potential mechanisms by which E4 exerts its larger AD risk effect (*i.e.*, in EA individuals), particularly directional transport along microtubules and golgi-ER trafficking.

We acknowledge limitations to our dataset and analyses. First, these were neurotypical, middle-age individuals, and whether donors would go on to develop AD is impossible to ascertain. Moreover, examining the molecular associations of E4 before the onset of clinical disease in a brain region where the first pathological changes may appear has the potential to uncover initiating pathological mechanisms of AD. However, E4 homozygosity has been described as a Mendelian form of pathological AD, at least in EA individuals^72^; to this point, our cohort has strong representation of E4 homozygotes (*n*=10; 2 EA). We also note that while we aimed to represent all combinations of EA/AA, E2/E4, and sex, our dataset does not contain EA E4 female donors, given the <20% frequency of the E4 allele and female underrepresentation in brain repositories^151^. These constraints likewise limited our power for sex-differential analyses and precluded analyses on risk factor interactions. Nonetheless, the representation of multiple epidemiologic risk factors is a strength overall. Second, despite the challenges interpreting our results due to the multicellular nature of spot-based SRT, (for example, astrocyte marker expression changes within the LC domain), alternative approaches would not readily reveal astrocytic changes specific to LC neuron local microenvironments. At the same time, we were able to infer astrocyte changes specific to LC, and associate spot-level NM to gene expression, highlighting strengths of this approach. The multicellular nature of SRT also impeded our ability to determine whether LC^NM-^spots contained putative NM-negative NE neurons, or simply were devoid of NM because they only contained noradrenergic cell processes (which do not generally contain NM). Finally, DE analyses did not examine the vascular domain, as PCA separated these samples primarily by their read depth (**Fig. S12**). However, LC plays an important role in blood-brain barrier (BBB) function^26,27^, and both *APOE* and sex show nominal associations with neurotypical BBB permeability^28^. Future studies at single cell and spatial resolutions will be needed to: decisively identify vasculature, NM-positive neurons, and NM-negative neurons in the LC; verify the cell types driving our DE observations; and determine whether NM-gene associations are LC-intrinsic or a function of cells surrounding LC neurons.

We have molecularly profiled the middle-aged human LC, representing the age range at which pTau accumulation, hyperexcitation, and degeneration begin to occur—events which are increasingly supported as etiologically important in AD. By collecting a large set of samples from this brain region, we further explore how this turning point in aging of the human LC is molecularly impacted by critical AD risk factors of sex, *APOE* haplotype, and ancestry. To facilitate community use of these data, we provide interactive web tools to visualize our SRT results and explore domain-level gene expression. Our data and findings promise to be a useful resource for developing therapeutics that aim to prevent the molecular changes that drive LC dysregulation, degeneration, and pTau propagation in AD.

## Supporting information

Legends for Supplemental Tables S1-S14 and Data S1-S5

## 4 | Acknowledgements, Funding, Authorship Contributions, and Data Availability

## 4.1 | Acknowledgements

We thank the LIBD neuropathology team, particularly James Tooke and Amy Deep-Soboslay for their work in sample curation and clinical characterization at LIBD. We thank Geo Pertea for maintaining and imputing the LIBD DNA genotype data. We thank the staff and physicians at the brain donation sites, and the generosity of the brain donors and their families, without whom this work would not be possible. We thank Dr. David Weinshenker for thoughtful comments on the analyses and the manuscript. Finally, we thank the families of Connie and Steve Lieber and Milton and Tamar Maltz for their generous support. Portions of some figures were created with Biorender.

## 4.4 | Data Access and Visualization

Raw data generated as a part of this study have been deposited on the Gene Expression Omnibus (GEO) with accession number GSE307866. Interactive web portals have been made available using *spatialLIBD*^90^ to visualize the spatial domain assignments and gene expression in the H&E stained tissue. Pseudobulk gene expression data is also provided in an *iSEE*^135^ app to allow users to browse domain expression patterns across domains, biological variables (sex, APOE haplotype, ancestry), and technical variables. The web applications to explore this data can be found on https://research.libd.org/LFF_spatial_LC/#data-visualization. **Supplemental Data S1-S5** and the processed data files (R objects) used to make the interactive apps can be found on https://research.libd.org/globus/ under the heading *jhpce#LFF_LC*.

## 4.5 | Code Availability

All code for processing, analyzing, and visualizing the SRT data are publicly available on GitHub at https://github.com/LieberInstitute/LFF_spatial_LC/. A freeze of the GitHub repository at the time of preprint is also made available through Zenodo: https://doi.org/10.5281/zenodo.17155605.

## 4.6 | Author Contributions

Conceived and Designed the Study: DRW, KRM, KM

Performed Experiments and Collected Data: HRD, SVB, RB, IDR, SM, AJS, AC, BAO, KDM, SHK, HA, AP

Software: RAM, LCT

Formal Analysis: BM, MT, SH, LHM, LCT

Data Curation: BM, HRD, LCT

Tissue Resources: JEK, TMH

Writing – original draft: BM, HRD, SVB, KM

Writing – review & editing: BM, HRD, SVB, TMH, DRW, SCH, SCP, KRM, KM

Visualization: BM, HRD, MT

Supervision: TMH, SCH, SCP, KRM, KM

Funding acquisition: BM, KRM, SCH, DRW, KM

Project Administration: DRW, KRM, KM

## 5 | Methods

### 5.1 | Postmortem human tissue samples and genotyping

Postmortem human brain tissue from the Lieber Institute for Brain Development (LIBD) Human Brain and Tissue Repository was collected through the following sites and protocols at the time of autopsy with informed consent from the legal next of kin: the Office of the Chief Medical Examiner of the State of Maryland, under the Maryland Department of Health IRB protocol #12–24, the County of Santa Clara Medical Examiner-Coroner Office in San Jose, CA, under WCG IRB protocol #20111080. Additional samples were consented through the National Institute of Mental Health Intramural Research Program under NIH protocol #90-M-0142, and were acquired by LIBD via material transfer agreement. Details of tissue acquisition, handling, processing, dissection, clinical characterization, diagnoses, neuropathological examinations, and quality control measures have been described previously^152^.

Routine AD pathology scoring was available for LIBD donors for donors over age 50 (**Table S1**), which confirmed little to no Aß or pTau pathology. Alzheimer’s neuropathology ratings include the Braak^153,154^ staging schema evaluating tau neurofibrillary tangle burden across several brain centers that have age-associated and Alzheimer’s positive tau pathology (neurofibrillary tangles and threads), and the CERAD (Consortium to Establish Ratings for Alzheimer’s Disease) scoring system^155^ as a measure of senile plaque burden (neuritic and diffuse). The ratings for Braak stage and numeric scores with CERAD scores for amyloid and neuritic plaques were updated from a combination of current rating systems below. An Alzheimer’s likelihood diagnosis is then performed based on the published consensus recommendations for postmortem diagnosis of AD as in prior publications^153–157^.

We obtained fresh-frozen transverse brainstem slabs containing the locus coeruleus (LC) from a cohort of middle-aged brain donors who were stratified by genetic risk for the *APOE2* (E2) versus *APOE4* (E4) alleles. The samples were collected from 33 neurotypical brain donors of both African American and European ancestry. The cohort contained 14 homozygote (E2/E2 and E4/E4) and 19 heterozygote (E2/E3 and E3/E4) donors from both ancestries (**Table S1**). For tissue block dissections, we identified the LC in transverse slabs of the brainstem at the level of the pons. The presence of the LC was determined through visual inspection of the slab, based on neuroanatomical landmarks and the neuromelanin pigmentation. Tissue blocks were dissected from the dorsal aspect of the pons, centered around the LC, using a razor blade, and stored at –80 °C until experimental use.

To determine genomic ancestry, genotype calling in all donors was performed using several Illumina BeadChip DNA microarrays. Using Plink v1.9^158^, we excluded variants with a minor allele frequency (MAF) less than 0.5%, a missingness rate of 5% or more, and a Hardy-Weinberg equilibrium (HWE) p<1x10^-5^. Genotype phasing and imputation were performed using the NHLBI NIH TOPMed imputation service with the GRCh38 TOPMed reference panel^159^. We finally filtered the imputed genotype data to keep only variants having MAF > 0.01. We inferred local and global ancestry in our study samples using *FLARE* (Fast Local Ancestry Estimation)^160^, which extends the Li and Stephens haplotype copying model within a hidden Markov framework^161^. Study sample genotypes were first phased and missing genotypes imputed with *Beagle*^162^. We used three reference populations from the 1000 Genomes Project: YRI (African), CHB (East Asian), and CEU (European). FLARE was run separately for each chromosome using the phased study genotypes, phased reference haplotypes, a GRCh38 genetic map, a minimum MAF threshold of 0.01 in the reference VCF (min-maf=0.01), and other default parameters.

### 5.2 | Tissue processing and anatomical validation

To anatomically validate LC inclusion on the tissue blocks, we evaluated cryosections using histology and multiplexed single-molecule fluorescence in situ hybridization (smFISH). For histology, we used a cryostat (Leica Biosystems) to cut 10 μm sections, which were adhered to glass slides, fixed with methanol and stained with hematoxylin and eosin (H&E) according to the manufacturer’s instructions (User guide CG000160 Rev C, 10X Genomics). Images of the H&E-stained slides were captured using a CS2 slide scanner (Leica Biosystems) equipped with a color camera, a 20x/0.75 NA objective and a 2x doubler for nominal 40x magnification. Brightfield images were saved as Tagged Image Files (.tif). We cut additional 10 μm cryosections, which were probed for the presence of norepinephrine neurons that mark the LC (*TH* and *DBH*), serotonergic neurons that mark the raphe nucleus (*TPH2*) and GABAergic neurons (*GAD2*) that mark the peri-LC. For smFISH experiments, we used RNAscope technology as described by Wang et al.^163^. The Fluorescent Multiplex Kit v.2 and the 4-plex Ancillary Kit (catalog no. 323100, 323120, ACD) were used according to the manufacturer’s instructions. Briefly, we fixed the 10 μm tissue sections (2–4 sections per donor) with 10% neutral buffered formalin solution (catalog no. HT501128, Sigma-Aldrich) for 30 min at room temperature. The sections were then dehydrated in a series of increasing ethanol concentrations (50%, 70%, 100%, and 100%), pretreated with hydrogen peroxide for 10 min at room temperature, and treated with protease IV for 30 min. Sections were incubated with these probes: *TH* (catalog no. 441651-C2, Advanced Cell Diagnostics); *DBH* (catalog no. 545791-C3, Advanced Cell Diagnostics); *TPH2* (catalog no. 471451-C1, Advanced Cell Diagnostics),; and *GAD2-ver2* (catalog no. 415691-C3). After probe labeling, sections were stored overnight in a 4× saline-sodium citrate buffer (catalog no. 10128-690, VWR). Following amplification steps (AMP1-3), probes were fluorescently labeled with Opal Dyes 520, 620, and 690 (catalog no. FP1487001KT, FP1488001KT, and FP1497001KT, Akoya Biosciences; all dyes were diluted to 1:500) and counter-stained with DAPI (4′,6-diamidino-2-phenylindole) to label cell nuclei for 20 sec at room temperature. After decanting, the sections were coverslipped using Fluoromount G mounting medium (catalog no. 0100-01; Southern Biotechnology). The coverslipped slides were stored in the dark for at least 24 hours before imaging. Slides were imaged using a Nikon AXR confocal microscope system powered by the NIS-Elements imaging software, utilizing a Nikon APO lambda D 20x / 0.80 objective and/or a Nikon APO lambda S 40x / 1.25 objective. For anatomical quality control, slides were imaged in a single plane using a Nikon Plan APO lambda D 2x / 0.1 objective (**Fig. S1**).

### 5.3 | SRT data generation

We used the Visium platform (10x Genomics) to generate SRT data. Visium Gene Expression Slides were cooled inside the cryostat, and 10 μm tissue sections were subsequently adhered to the slides. The tissue sections were fixed with methanol and stained with H&E according to the manufacturer’s instructions (User Guide CG000160 Rev C). Following H&E staining and image acquisition, the slides were processed for the Visium assay according to the manufacturer’s instructions (Visium Gene Expression Protocol User Guide CG000239, Rev D). Briefly, the workflow included permeabilizing the tissue to allow access to mRNA, followed by reverse transcription, cDNA removal from the slide, and library construction. Tissue permeabilization experiments conducted on a single LC sample identified an optimal permeabilization time of 18 min, which was used for all sections across donors. To maximize capture efficiency, tissue blocks were scored so that two adjacent tissue sections from the same donor could be placed onto a single Visium capture area, resulting in a total of 43 Visium capture areas.

Following quality control, libraries were sequenced on the NovaSeq 6000 Illumina platform at the Johns Hopkins Single Cell Transcriptomics core, following the manufacturer’s instructions with a minimum sequencing depth of 60,000 reads per spot. Samples were sequenced to a median depth of 263,316,447 reads resulting in a median 76,189 reads, 1,152 unique molecular identifiers (UMIs), and 526 median genes per spot.

### 5.4 | Raw SRT data processing

Sample slide images were first processed using *VistoSeg*^164^. *VistoSeg* was used to divide the Visium sample slides into individual capture areas using the *VistoSeg::splitSlide()* function. This function takes one large image from the entire slide and separates it into individual capture areas, each capture area on the slide labelled as A1, B1, C1 and D1. These individual capture area images were used as one of the inputs for SpaceRanger (10x Genomics).

Individual capture area images from *VistoSeg* were spatially aligned to Visium Slide capture areas using the *Loupe Browser* (version 7.0.0, 10x Genomics). This alignment enabled precise correspondence between histological features and individual capture spots on the slide. Following alignment, raw sequencing data were processed with *SpaceRanger* software (Version 3.0.0, 10x Genomics). The workflow incorporated the *Loupe Browser*’s JSON file, aligned tissue images, and associated FASTQ files from sequencing to generate spot-by-spot spatial feature counts for each sample. The resulting *SpaceRanger* output metrics were subsequently imported into *R* using the *spatialLIBD* Bioconductor package^90^. *SpatialExperiment* objects were constructed for downstream spatial transcriptomics analyses, facilitating integration with established workflows and reproducible data exploration^165^.

### 5.5 | H&E image processing and quantification of NM signal

#### 5.5.1 | Segmentation

High-resolution H&E images (**Fig. S3A**) from all samples were processed using *VistoSeg*^164^, with the *VistoSeg::VNS()* function customised to use a cluster resolution of *k*=2 instead of the default *k*=5. This strategy resulted in one cluster capturing a white-to-black gradient, successfully isolating neuromelanin (NM) pigment (**Fig. S3B**). We then converted the NM-associated color cluster to grayscale, with intensity values ranging from 0 to 1. Intensity distributions across capture areas were examined, revealing consistent patterns across samples within the 0–0.6 range (**Fig. S3C**). A threshold of 0.5 was applied to extract the dark NM pixels, while a lower threshold of 0.2 was used to exclude super-black pixels resulting from debris or other artifacts. Some noise persisted at the whole-tissue section level, as artifacts such as tissue folds, curls, or tears often resembled NM pixels in color, making them difficult to exclude using size or intensity thresholds alone (**Fig. S3D**). To address this, image processing like morphological operations—specifically image opening followed by closing—were applied in *MATLAB*. The opening operation removed small, noisy elements misidentified as NM, while the closing step recovered fragmented true NM signals. This combined approach substantially reduced background noise that size filtering failed to eliminate (**Fig. S3E**). Documentation on *MATLAB* morphological operations can be found at^166^.

#### 5.5.2 | Quantification and Spot Classification

The refined segmentations were processed using the *VistoSeg::CountNuclei()* function to extract spot-level NM metrics and classify spots^164^. For each SRT spot, the resulting output included: (1) the proportion of the spot area covered by segmented NM pixels, (2) the mean intensity of those pixels, and (3) the number of segmented NM regions. Based on the distribution of NM coverage across samples (**Fig. S3F**), a threshold of 0.3% NM area per spot was used to classify spots as NM+ (≥ 0.3%) or NM–(< 0.3%) (**Fig. 1C**). Small, distinct segmented speckles after post-processing (**Fig. S3E**) often located near tissue edges or tears, contributed to low-coverage (< 0.3%) spots that were considered background noise.

#### 5.5.3 | Calculation of Tissue-Level LC NM Score

Once LC domain spots were annotated by clustering (*Methods 4.6.3*), a polygon was drawn around the LC region on each tissue section within the capture area. The mean intensity of segmented NM pixels within this polygon was then calculated and exported as the tissue-level NM score for each tissue section.

### 5.6 SRT data analysis

All analyses were performed using R 4.4 except where noted. Data was analyzed as a *SpatialExperiment*^165^. *spatialLIBD*^90^ and *escheR*^167^ were used to plot spotwise gene expression. The *ggplot2*^168^ package was used to generate general-purpose visuals such as scatter plots and heatmaps. Extended color palettes were generated using *Polychrome*^169^. Key R packages and versions used were as follows: *spatialLIBD* 1.18.0, *SpatialExperiment* 1.16.0, *RcppML* 0.3.7, *scater* 1.34.0, *scran* 1.34.0, *scuttle* 1.16.0, *scry* 1.16.0, *nnSVG* 1.10.0, *harmony* 1.2.3, *bluster* 1.16.0, *spacexr* 2.2.1, *WeberDivechaLCdata* 1.8.0, *edgeR* 4.4.2, *limma* 3.62.2, *SpotSweeper* 1.0.2, and *fgsea* 1.32.4.

#### 5.6.1 | Sample QC and count normalization

First, we removed the outermost spots surrounding a tissue sample, *i.e.* those along its edges, using a custom script (see *Code Availability*). Subsequently, we used *SpotSweeper*^170^ to detect spots that were outliers (relative to nearby spots) with high mitochondrial read percentage, low total unique molecular identifiers (UMIs), and/or low number of unique genes detected, and removed all such spots. A hard filter for mitochondrial read content was not employed because mitochondrial counts are correlated with known biology in the LC^91^ and in SRT approaches generally^75,171^, and differed across tissue sections. Many remaining spots after this filtering step still had very low content (10th percentile for total UMIs < 100 and for unique genes < 40); this was unsurprising since much of the tissue surrounding the LC is myelinated white matter (WM) tracts, which are RNA-poor relative to gray matter^91,172^. Therefore, our final filtering step removed spots in the bottom 1st percentile for UMI and/or unique gene counts (<24 or <18, respectively). One Visium capture area contains 4,992 spots; in this dataset, tissue covered 3,481±409 (mean ± standard deviation (SD)) spots before filtering and 3,081±511 after. The retained spots contained 1,925 ± 2,521 UMIs (mean±SD) across 855 ± 909 unique genes, constituting a final dataset of 132,484 spots and 29,877 genes with at least one count. Additional post-filtering metrics are included in **Table S1**. Log-normalized counts were then calculated for all remaining spots based on the number of UMI counts per spot (library sizes) using *scater*::*computeLibraryFactors*() followed by *scater*::*logNormCounts()*, both with default parameters^173^.

#### 5.6.2 | In silico separation of tissue sections into SRT samples

With one exception, all SRT capture areas contained two consecutive tissue sections from an individual donor. To leverage this experimentally-designed replication, we separated each capture area into two tissue sections (“samples”) by making interactively annotating tissue and non-tissue spots as one of two ‘clusters’ using *spatialLIBD*^90^. The spot identifiers were then matched back to their capture area to visualize which sample they had been assigned to, and manually refined as necessary. These annotations were used to then append a slice identifier (1 or 2) to the end of each spot identifier and to divide capture areas into multiple samples. As all spot IDs within a capture area are already unique, the section and capture identifiers could be interchanged to allow for analyses at the levels of capture area or individual tissue sections. We use the term “sample” to denote *one* tissue section from a capture area, regardless of how many tissue sections that capture area contained in total. For tissue section-level applications, log-normalized counts were recalculated^173^ considering each section as its own sample.

#### 5.6.3 | Feature selection, dimensionality reduction, preliminary clustering, and clustering assessment

Multiple approaches to feature selection were initially used to identify informative features for dimensionality reduction. Using the log_2_-normalized counts, we identified highly-variable genes (HVGs) in *scran*^174^. Given the large quantity of low-read spots that we had noted during QC, we also utilized a raw-count based measure of gene variance across spots to define high deviance genes (HDGs)^76^. Finally, we identified spatially-variable genes (SVGs) using *nnSVG*^77^ per sample (single tissue section), considering genes with ≥ 2 counts in ≥ 2% of spots for a given sample. From each sample, gene identifiers and SVG ranking were collected for genes with nominally significant spatial variation. To get dataset-level SVGs, 762 genes with nominally significant spatial variation in ≥43 samples (*i.e.*, >50% of the 85 total samples) were retained, and their rank across the nominally significant samples averaged to get a data-wide rank. Several feature sets were then generated, capped at 10% of the number of genes analyzable using the HVG approach, or 1,397 genes. We took the top 1,397 HDGs, all 762 SVGs, or 1,397 HVGs as single-approach feature sets, as well as mixtures of these by combining 0.25, 0.5, or 0.75 * 1,397 genes from HDG Fig or HVG with SVGs. After removing duplicate genes, these sets ranged in size from 762-1,397 genes. For example, the presented feature set of 765 genes, “25% HDGs” and “75% SVGs”, contained all 762 SVGs and the 3 top 349 HDGs that were not also SVGs.

For each feature set, we then performed dimensionality reduction using standard PCA and, separately, a normalization-free approach termed *glmPCA*^76^. Each dimensionality reduction was then batch-corrected across samples by inputting all 50 principal components (PCs) into *Harmony*^78^ with lambda set to null to use data-driven initialization of the lambda (correction strength) parameter. The output from *Harmony* was used as input to Louvain clustering^79^ at resolutions of 0.5, 1.0, and 2.0. Putative noradrenergic cluster(s) were identified based on marker analyses of each clustering, identifying clusters enriched for *DBH* > 2-fold vs. all other clusters at a statistically significant level (FDR<0.05).

We assessed the internal validity of LC-NE identification from each clustering approach by manually defining LC spots under low- and high-stringency conditions. The high-stringency definition required a spot to have ≥1 count of *DBH* and ≥1 count of either *TH* or *SLC6A2*, with a total of ≥5 counts, as well as ≥2 counts of additional markers (*PHOX2A*, *PHOX2B*, *DDC*, *SLC18A2*). The low-stringency definition required ≥1 count of *DBH* and ≥1 count of either *TH* or *SLC6A2* along with at least 1 count of any of the additional markers. We then calculated the Spearman correlation of expression in LC pseudobulked by sample or by donor when defining it from a clustering approach (collapsing multiple LC clusters into a single one if present) vs. by assigning one of the manual definitions above. Surprisingly, most samples showed consistently strong concordance between expression profiles of clustered LC and high-stringency manually-defined LC regardless of feature set or dimensionality reduction approach. *glmPCA*-defined clusters performed slightly worse for a given feature set and clustering resolution and were therefore not considered further. Of the remaining approaches, samples that showed less consistent agreement with manual LC definitions were used to prioritize the most robust feature and clustering parameter sets. The final clustering approach was selected based on visualization of cluster labels superimposed on tissue histology to assess whether LC cluster labels were being assigned well outside of areas with plainly visible neuromelanin. The average Spearman correlation between each sample’s manually defined LC spots and LC defined with the final parameter set (765 genes, “25% HDGs” and “75% SVGs”, with standard PCA and Louvain clustering at resolution 1.0) was >0.95 for both high- and low-stringency manual definitions. We refer to the final, annotated clusters reported in the text as “domains”.

#### 5.6.4 | Domain annotation

The unsupervised clustering described above yielded 9 domains. In reassessing sample anatomy alongside the domain annotations, we could not definitively confirm the anatomy to be in an LC-containing brain area for three donors: Br5517, Br5276, and Br5712. All analyses reported henceforward were thus performed using the remaining *n*=30 donors. Domains were annotated based on their top-enriched genes identified using one-vs-all markers analysis implemented in *spatialLIBD*^90^ using samples from the 30 donors retained in downstream analyses. The LC domain was readily defined by enrichment of canonical markers including *SLC6A2* and *DBH*. To split LC spots into subdomains of NM-positive and NM-negative (LC^NM+^ and LC^NM-^, respectively), we applied the spot classifications generated in Section 4.5.2, which were derived from the NM segmentation metrics (**Fig. S3F**). Specifically, LC^NM+^ spots were defined as those within the LC domain exhibiting ≥0.3% of their area covered by segmented NM pixels, while the remaining LC domain spots are designated LC^NM-^.

Two neuronal domains showed unexpected sets of marker genes: both were significantly enriched for serotonin transporter *SLC6A4*, acetylcholinesterase (*ACHE*), as well as the serotonin receptor *HTR2C*. We leveraged recent brain-wide mouse^82^ and human^83^ single-cell/nucleus datasets as well as previous cell type-enriched expression profiles of mouse serotonergic neurons^80,81^, to determine that *HTR2C* and *SLC6A4* are seldom co-expressed. Further, these domains were both enriched for excitatory marker *SLC17A6*, yet also for the neuropeptide *ADCYAP1*. Given the perplexing set of marker genes identified, these two domains were thus denoted using two of their distinctive top marker genes (“*ACHE*, *SERT*…”).

Three oligodendrocyte (oligo) domains were readily distinguished by top fold-enrichments of *KLK6* and genes encoding protein components of myelin (e.g., *MOBP*, *MAG*, *MOG*). Astrocyte domains were distinguished by *FGFR3*, *AQP4*, and *APOE*. The remaining domain was very heavily enriched in hemoglobins (*HBA*, *HBB*) and clearly corresponded to blood vessels when plotted onto tissue samples and was thus labeled Vascular.

To identify features differentiating between the two *ACHE-SERT* domains, the three oligodendrocyte marker-enriched domains, or the two astrocyte marker-enriched domains, we re-ran marker analysis as described above to compare each domain individually to other(s) of the major type (e.g., only among oligodendrocyte marker-enriched domains; **Table S4**). The relative marker enrichments were used to then annotate additional cell type(s) especially well-represented in one domain.

Direct comparison of the astrocyte-rich domains revealed one domain was comparatively enriched for markers of oligodendrocytes, including *MBP* and *MOBP*, suggesting such spots were astrocyte-oligodendrocyte mixtures with astrocytic transcripts as the predominant content (Astro-Oligo). The second domain was comparatively enriched for additional astrocytic markers such as *SOX9*, *AEBP1*, and *PLTP*. Thus, we annotated this domain as primarily astrocytic (Astro; **Table S4**). Three oligodendrocytic domains were enriched in two or more genes encoding myelin components (*MBP*, *MOBP*, *MOG*, *MAG*; **Fig. 2C**). One-vs-others marker analyses (**Table S4**) among these domains revealed one domain (Oligo) to be comparatively enriched for canonical markers such as *MOG* and *KLK6*, suggesting this domain had the purest oligodendrocyte representation. A second domain (Oligo-Astro) was comparatively enriched for astrocytic and endothelial markers (*e.g. VIM* and *CLDN5*), while the third domain (Oligo and Other Cell Types Mixed) was enriched for both astrocytic genes (e.g., *SLC1A2*, *FGFR3*) and neuronal genes (e.g., *CAMK2B*; **Table S4**). In the *ACHE-SERT* domains, one was comparatively enriched for several additional neuropeptides including *SST*—which would intuitively suggest interneurons—while the other was comparatively enriched for oligodendrocyte and LC markers. Thus, the final *ACHE*-*SERT* labels were “*ACHE*-*SERT* Neuropeptides Mixed” and “*ACHE*, *SERT*, LC, WM Mixed”, with “WM” denoting white matter and “mixed” denoting mixed cell type identities in the domain.

To determine whether there were characterized human cell types underlying the transcriptomic profile of *ACHE*-*SERT* spots, we used previously published snRNA-seq profiling of adult human LC^91^ as reference data for spot deconvolution implemented in *RCTD*^92^. Our previous snRNA-seq data includes unannotated granular cluster labels and their broader groupings into major cell types (excitatory neurons, inhibitory neurons, etc), although each granular cluster was placed into one of the named merged clusters. Therefore, we concatenated each unannotated granular label (numbers from 0-29) with the merged label to which it was assigned for interpretability. We then ran RCTD in “multi” mode, allowing for up to 4 cell types to be contributing to a spot’s transcriptome, as well as “full” mode, which has no cap on the number of cell types represented in a spot. Other run parameters specified during data setup with *create.RCTD* were *UMI_max=Inf*, *UMI_min=50*, *DOUBLET_THRESHOLD*=0, and *counts_MIN*=10. We then extracted the weights (estimated proportion of spot transcripts attributable to a cluster in the reference data) assigned to each spot from “multi” and “full” modes, which demonstrated high weights for a particular granular cluster, #4 (“inhibitory_4”). To determine the distinguishing feature of the inhibitory_4 population in the snRNA-seq data, we performed marker analysis in *spatialLIBD*^90^ as described above, comparing inhibitory_4 against the rest of cells annotated as inhibitory under “label_merged”.

#### 5.6.5 | Intra-LC spatially-variable gene analysis

In order to determine whether there was spatially heterogeneous expression of genes within the LC domain specifically, we again leveraged *nnSVG*^77^, analyzing only LC domain spots in each sample (single tissue section), considering genes with ≥ 2 counts in ≥4 LC spots of a given sample. *n*=68 samples contained adequate LC spots to perform SVG analysis. Per *nnSVG*^77^ documentation, we only used the BRISC-AMMD spatial-ordering algorithm for samples containing 65 or more LC spots, and used the argument *order=”sum_coords”* for all other cases. We then aggregated the results from these 68 samples and visualized genes that were nominally significant SVGs in ≥20 samples (guaranteeing the result came from ≥5 donors). We additionally tabulated the gene ranks from each sample and calculated the mean rank across those samples in which a gene was a nominally significant SVG (**Table S2**).

#### 5.6.6 | NMF of LC gene expression

To decompose LC gene expression signals into constituent expression programs, we applied non-negative matrix factorization (NMF) only to spots included in the LC domain. Specifically, *RcppML*^118^ was used to identify 160 factors. 160 was chosen as an estimate calculated as number of distinct AD risk factor groupings: 2 sexes by 4 *APOE* haplotype pairs by 2 predominant ancestries, assuming 10 granular factors unique to a given grouping, yields 2*4*2*10=160). NMF was performed using data filtered to genes with at least 56 counts (15.2k genes) across LC spots of samples from the *n*=30 donors retained for downstream analyses. (As one of these retained donors did not have any spots containing LC, *n*=29 donors for this analysis).

We then looked for genes that were specific to one or a few factors, and factors that were enriched for such genes. We defined “factor-specific” genes as those with a factor loading above 1.5 x the median absolute deviation of loadings for that gene across all factors that only achieved a loading of this degree in 3 or fewer factors total. Gene sets were then defined for each factor that contained at least 25 such genes (a total of 15 factors and 1,563 unique genes; some factors shared genes, for a total of 2,300 factor-gene pairings).

Each set of factor-specific genes was tested in the *enrichR*^105,175^ package, using all genes included in the NMF calculation as the background set, in order to attribute molecular and cellular functions to factor-specific gene sets.

To determine the cell type expression patterns most consistent with individual NMF factors, we projected the NMF matrix into our prior human snRNA-seq dataset^91^ using *RcppML*^118^. For each snRNA-seq cluster, we then calculated the proportion of nuclei that were given a non-zero weight for a given NMF factor and the scaled average weight for that factor across all nuclei of the cluster, allowing for visualization of factor-cell type relationships as in **Fig. 3**. Additionally, we projected our NMF matrices into recent snRNA-seq from the mouse LC^87^ using all genes with 1-1 human-mouse orthologs detected in both datasets. This procedure was performed using the authors’ broader label set, consisting of one LC/*Dbh+* cluster.

#### 5.6.7 | Differential expression analyses and gene set enrichment analyses (GSEA)

To examine differences in gene expression, we pseudobulked gene counts by domain within each tissue section, and removed pseudobulks representing samples with fewer than 10 spots from a domain. For example, if a sample had 7 Astro spots, the Astro pseudobulk from that sample was removed. We then examined dimensionality reduction of these pseudobulk profiles. Four domain profiles distributed in space mostly as a function of their library size (i.e., gene count)--as these were low-read depth domains, they were excluded from differential expression analyses. The domains not included for DE were the Vascular, Oligo, Astrocyte-Oligo Mixed, and Oligo-Other Types Mixed.

The remaining pseudobulk profiles from the other 5 domains—LC, two *ACHE*-*SERT* domains, Astro, and Oligo-Astro mixed—were combined into one large gene expression matrix. We chose to model all domains jointly as our findings suggested that most spots were, in reality, mixtures of cell types, such that useful transcriptomic information for a given cell type is contained within several spot domains. Models controlling for Age, Sex, *APOE* (E2 or E4 carrier status), genomic ancestry proportion as a continuous value 0-1 (1=100% YRI/AA; 0=100% CEU/EA), and donor as a blocking factor (to account for multiple tissue sections per donor) were run separately for each AD variable of interest. Each model used a collapsed variable encompassing domain label of each pseudobulk with the AD variable of interest. For example, sex DE used a variable column encoding sex and label (“LC_male”, “LC_female”, “Astro_male”, “Astro_female”, …) and assessing contrasts (“LC_male -LC_female”, “Astro_male - Astro_female”) between the sexes for each pseudobulked domain. In the case of *APOE* haplotypes, we focus primarily on contrasts between all carriers of E4 and of E2, but did additionally perform contrasts between haplotype subgroups (e.g., E4 homozygotes vs. E4 heterozygotes). For ancestry-differential expression and ancestry-stratified *APOE* haplotype analyses, we categorized donors based on predominant genomic ancestry (≥75% or ≤25% YRI); 4 donors fell between these thresholds and were thus removed from the ancestry-differential analysis (**Table S1**). For each DE analysis, lowly expressed genes were first identified and removed using *filterByExpr* in *edgeR*^176^, providing the counts and DE model being tested as input. We then modeled gene expression differences in each domain using *voomLmFit* from the *edgeR* package^176^, with sample.weights set to TRUE to leverage its *voomWithQualityWeights*^177^–like sample weighting, and *eBayes* with *robust*=TRUE^178^. For DE analysis between LC^NM+^ and LC^NM-^, we used the same procedure and controlled for the same cohort variables, but only considered pseudobulks of the LC subdomains.

DE models did not control for RNA integrity numbers (RINs), as these were collected from cortices of the donors, which undergo a separate freezing and slabbing process compared to hindbrain, and thus are not representative of potential RNA degradation in hindbrain samples during initial tissue banking.

To functionally interpret DE findings, we performed Gene Set Enrichment Analysis (GSEA)^102,126^ via the R package *fgsea*^179^ to analyze DEG enrichment in sets from MSigDB^102–104^, including “hallmark” sets, sets by chromosomal region; curated sets from literature and databases (e.g., KEGG pathways); inferred microRNA (miRNA) / transcription factor (TF) target genes; gene ontologies (e.g., GO); and cell type markers. (These genesets are termed “h.all”, “c1.all”, “c2.all”, “c3.all”, “c5.all”, and “c8.all”, respectively). In separate GSEA analyses, we leveraged our previously aggregated^99^, tissue-nonspecific sets of putative TF target genes ascertained from model organisms and human cell/tissue types as indexed by *Enrichr*. Briefly, these sets included ChEA 2022^180^, TRRUST^181^, literature supplement-mined sets from Rummagene^106^, and those curated in the *Enrichr*^105^ tool itself (Gene Expression Omnibus-mined results of TF perturbation and knockout experiments and TF protein-protein interactions). For TF-target GSEA analyses, the results were subsequently filtered to TFs with at least weak expression in the domain analyzed, considering that TF mRNAs are generally lowly expressed^182^. This filtering examined spots from the analyzed domain, retaining TFs that were detected in at least 4% of domain spots in at least 12 samples (half the size of the smallest DE group tested for any domain, which was LC in EA donors), *or* with an of average across all samples of >2% domain spots containing the TF. In the case of TF-target GSEA on our DE results between NM+ and NM-LC spots, we used lenient filters to account for the relatively small number of spots available, filtering to TFs expressed in at least 1% of the LC subdomain’s spots in at least 25% of samples and a mean % of expressing spots >0.75% in the subdomain across all samples.

Gene sets analyzed were limited to those containing 15-500 genes for MSigDb and TF-target analyses. Each analysis used the signed DE *t*-statistics from one comparison for one domain (e.g., sex in the LC domain) as the ranked list for analysis. To retrieve enrichment statistics from every gene set for both directions of DE, we explicitly ran analyses to test enrichment at the top (positive logFC; *t>0*) and bottom (negative logFC; *t*<0) of the rank lists, as *fgsea* testing on both directions simultaneously will otherwise return data for the effect direction with greater enrichment, even if both directions are significant. For each DE comparison in a domain, a second GSEA analysis was also performed using the absolute value of DE *t*-statistics to identify gene sets enriched for DE as a general phenomenon (*i.e.*, regardless of the direction of effect). We used the *fgsea* parameter *eps=*0.0 to obtain a *p*-values estimate for all tests and specified 50,000 permutations for initial *p*-value estimations.

#### 5.6.8 | Gene-regulatory network (GRN) inference using SCORPION

Our analyses of LC heterogeneity included asking whether distinct transcriptional-regulatory patterns may distinguish LC^NM+^ and LC^NM-^ subdomains. To infer gene-regulatory networks without functional-omics data from the LC *per se*, we leveraged *SCORPION*^112^, a tool that adapts the PANDA^113^ message-passing algorithm for inferring gene regulatory networks in high-dimensionality transcriptomic data. We followed the practice from the original publication describing *SCORPION*^112^ of using reference datasets of TF-target gene pairs^114^ and TF-TF protein interactions from STRINGdb^183^ alongside our gene expression data. We subsetted to LC^NM+^ and LC^NM-^ spots and filtered to those genes that passed low-expression filtering in the DE test comparing these subdomains, and subsequently filtered the reference datasets to the same genes. We then separately ran *SCORPION* on each subdomain to obtain individual networks for comparison. *SCORPION* performs dimensionality reduction using PCA on user-provided expression data to create small pseudobulks from the most closely related spots or cells to address read sparsity. We specified *nPC*=20 (vs. default 25) as we were analyzing only one subdomain and set *gammaValue=6* such that 6 spots would constitute each pseudobulk used for GRN construction. We used *assocMethod=”Spearman”*, keeping other arguments set to their default values. We then obtained the resulting TF-gene regulatory networks, which have *z*-scored edge values representing the likelihood of a regulatory relationship–enhancing or repressing–between one TF and transcription of one gene (thus, large positive edge weights indicate strong likelihood of a gene’s regulation by a TF, even if the TF is acting as a transcriptional repressor).

To compare the LC^NM+^ and LC^NM-^ GRNs, we used tutorials from the developers of PANDA (https://netzoo.github.io/netZooR/ and https://github.com/kuijjerlab/multi-omics-course/blob/main/multi-omics_workshop_full_code.ipynb) to identified those edges with a high weight and a large difference in weights between the two networks. Specifically, the base R function *pnorm()* was used to identify edges in each GRN with >75% probability of representing ‘true’ interactions, and specific edges were defined as those for which pnorm(edge^NM+^) * pnorm(edge^NM+^ - edge^NM-^) was also >0.75. This yielded a network of ∼1,669 TF-gene relationships specific to LC^NM+^ or LC^NM-^. We also identified TFs with mostly LC^NM+/-^-specific edges, namely those TFs with ≥80% of edges specific to one LC^NM^ subdomain network. The 1,669 specific network edges, and an extended table including those edges plus all edges involving an aforementioned ≥80%-specific TF are in **Table S8**, and a full table of GRN output with all edge weights from LC^NM+^ and LC^NM-^ is provided in **Data S3**.

#### 5.6.9 | NM-associated gene expression and enrichment

In order to associate gene expression with local NM content, normalized log-counts were obtained for the 3,145 spots defined as NM-positive (see *Methods 4.5*) in 73 tissue sections from 29 donors (33 to 278 spots per donor, 7 to 152 per tissue section). We then modeled spot-level intensity of NM (the mean pixel intensity of segmented NM pixels within a spot; lower values signifying darker areas and thus stronger pigmentation) and proportion of spot pixels segmented as NM-positive as a function of gene expression. For each of the 23,744 genes detected in at least one spot, linear mixed models (LMMs) were analyzed in R using *lme4*::*lmer()*^184^ with the model *NMvariable* ∼ *logcounts + E4carrier + Sex + Age + Ancestry + (1|Donor) + (1|Donor:captureArea) + (1|Donor:captureArea:tissueSection)*, where *E4carrier* encoded whether the donor was an E4 carrier or not (with all non-E4 carriers being E2 carriers by design), age a continuous measure in years, and ancestry being estimated proportion YRI genomic ancestry. The aforementioned model also contained random intercept terms to account for nested levels of repeated measurements, specifically the fact that for every donor, two tissue sections (samples) were contained within each SRT capture area (that is, stained and sequenced together during data collection); and that, for some donors, two capture areas (four samples) were collected on separate SRT slides.

For each NM metric-by-gene model, the REML model coefficients were tested for significance and extracted into a table using *car::Anova()*. A log-ratio test was also performed comparing *non*-REML versions of the model specified above and a second without a term for the gene’s expression. The results for all fixed effects from each gene model for each NM metric were then tabulated. For each gene, the unadjusted *Anova* p-values were FDR-corrected and Bonferroni-adjusted using the number of genes successfully modeled (≤ 23,744).

To examine whether NM-associated genes were enriched in functional pathways, ontologies, or cell/disease marker genes, the *enrichR* package was utilized in R to query dozens of gene set libraries simultaneously^105,175^. We performed separate analyses for NM pixel proportion-associated genes and NM intensity-associated genes, treating all genes analyzed as the background set with query genes being those significantly NM-associated (FDR<0.05 or Bonferroni<0.05, separate analyses) with positive coefficients only, negative coefficients only, or their union (total of 6 significance threshold-effect direction combinations per NM metric). The main findings report results considering Bonferroni-significant genes associated with NM intensity in either direction, but we provide tables of top enrichment results for each NM metric at each significance threshold and effect direction (**Table S14**).

## Supplemental Table Legends

**Table S1. Donor demographics, demographic/analysis groups, and RNA-seq metrics from the filtered Visium data. A)** Demographic variables and, where available, tau and Aß pathology ratings (*Methods*) for each donor. Ancestry is assigned based on predominant (≥50%) global genomic ancestry estimates (*Methods*). **B)** *N* and demographics for the donors included in DE and other downstream analyses (up to 30). Subgroupings are provided by sex, predominant ancestry, and *APOE* haplotype. Only those donors considered in DE and other downstream analyses (up to 30) are considered. **C)** Demographic and other variables frequently considered in RNA-seq analyses by Visium capture area. RNA integrity number (RIN) are listed for cortical tissue from the same donors (although dissection and freezing of brainstem is performed separately from cortex during tissue banking). Also provided are means and medians of spot-level RNA-seq metrics per capture area after removing low-content and local outlier spots.

**Table S2**. **Spatially variable genes (SVGs) and high-binomial deviance gene (HDGs) analyses; spatial registration**^88–90^ **with human and mouse LC snRNA-seq data.** SVG sheets include a summary sheet of analysis, listing gene ranks for nominally significant genes in each sample, and separately including *nnSVG*^77^ output for all genes analyzed in each sample. A third sheet provides full HDG analysis^76^ results used to identify HDGs. The first three sheets contain a column to indicate genes used for clustering. Spatial registration results for the 9 reported domains with human^91^ LC and mouse^87^ RNA-seq datasets, calculated over a range of 10 to >1000 top-ranked marker genes for each reference cluster/domain.

**Table S3**. **Clustering QC metrics for the assigned domain labels; domain proportions by donor and tissue section.** The first sheet contains the spot identifier, assigned label, 3 columns of results from cluster purity analysis, a column for silhouette width, and a column for root mean-square deviation (one value per domain). The remaining sheets detail the proportion of spots assigned each domain by brain donor or by tissue section.

**Table S4**. **Domain marker gene analysis and domain comparisons among *ACHE*-*SERT*, Oligo, or Astro domains.** The first sheet provides the full results of one-vs-all marker analyses for the 9 reported domains. The remaining sheets include analogous results when only comparing among Astro domains, among Oligo domains, or among *ACHE-SERT* domains.

**Table S5**. **snRNA-seq marker analysis of *ACHE*-*SERT* domain-enriched inhibitory subcluster 4.** Marker analysis table for the inhibitory cluster from^91^ predicted to be a large source of expression in *ACHE*-*SERT* domains by *RCTD*^92^. Also see **Fig. S10**.

**Table S6**. **nnSVG results from analysis restricted to the LC domain**. A summary sheet of the analysis, lists ranks for nominally significant genes in the LC domain of each sample, and separately including *nnSVG*^77^ output for all genes analyzed in each sample. Note that some LC-containing samples with insufficient spots to determine spatial variation were not analyzed.

**Table S7**. **DE and GSEA analyses comparing LC^NM+^ and LC^NM-^.** Sheet with DE results, where positive logFC signifies greater expression in LC^NM+^ relative to LC^NM-^. A second sheet is concatenated results from two GSEA analyses (one examining MSigDB gene sets, one examining sets of TF-regulated genes). TF-target results were subsetted to those where the TF was at least marginally expressed in LC (see *Methods*). Additional information is included in the GSEA sheet to aid in table interpretation.

**Table S8**. *SCORPION* GRN inferred LC^NM+^/LC^NM-^-specific gene-regulatory relationships and TFs biased toward LC^NM+^/LC^NM-^-specific activity. A table of the ∼1,669 GRN edges (TF-gene pairs) deemed LC^NM+^/LC^NM-^-specific (see *Methods*). A second sheet contains the 1,669 aforementioned edges plus all edges involving a TF with LC^NM+^/LC^NM-^-specific activity. The third sheet lists these TFs, which are those with ≥80% of edges specific to either subdomain being LC^NM+^ or LC^NM-^-specific; that is, *# subdomain-specific edges* / *(# of LC*^NM+^ *specific edges + # of LC*^NM-^ *specific edges)*>0.8.

**Table S9**. **Genes unique to one-few LC NMF factors and Enrichr analysis of NMF factor-specific genes**. Genes identified as uniquely upweighted in 1-3 NMF factors are listed, with a row for each factor-gene pair. Enrichr results with an adjusted *p*-value<0.05 are provided alongside details to aid in table interpretation. The background gene list used for enrichment testing in this analysis is also included.

**Table S10**. **DE results from 5 domains compared between sexes, E4 and E2 controlling for ancestry, ancestry-stratified E4-E2 comparisons, and between predominant ancestries.** Each sheet’s title indicates the comparison, with the group in which a positive logFC signifies upregulation listed first (e.g., ‘male-female’). A legend sheet is also included.

**Table S11**. **MSigDB and TF-target GSEA analyses on sex-DE and haplotype-DE.** MSigDB GSEA results significant at an FDR<0.05 are included for each DE analysis. For TF-target analyses, results were subsetted to those where the corresponding TF of interest was expressed in the analyzed domain (see *Methods*). Sheets include a top row with column definitions and sign conventions to facilitate interpretation.

**Table S12**. **Ancestry-haplotype DE enrichments in MSigDB gene sets.** The first sheet lists those GSEA results that were significant (FDR<0.05) from E4-E2 analysis in a domain for only one ancestry. The remaining sheets provide all FDR<0.05 GSEA results from ancestry-specific haplotype-DE (AA E4-E2 and EA E4-E2) and single-ancestry haplotype-DE (E4 AA-EA and E2 AA-EA).

**Table S13**. **Results from linear mixed models of spot-level NM intensity or spot NM pixel proportion against spot-level gene expression.** Sheets provide results of the reported NM intensity modeling as well as modeling of NM pixel proportion. For each gene, rows are included with the coefficient and raw *p*-value for each fixed effect (sex, E4/E2 carrier, proportion African genomic ancestry, age, and gene expression) and for the log ratio test statistic comparing models of the NM metric with and without gene expression. For gene coefficients, the FDR- and Bonferroni-corrected p values are also listed. Sheets include additional details to aid in interpretation.

**Table S14**. ***Enrichr* analyses of NM intensity- or proportion-associated genes.** Top results from 6 *Enrichr* analyses per NM metric modeled: two significance cutoffs for NM association (FDR<0.05, Bonferroni *p*<0.05) and three bins by sign of association (positive, negative, all). *Enrichr* reports the rank of each test within a library of gene sets^105,175^; up to 20 significant enrichments are listed for each *Enrichr* library and input set of NM-associated genes. A sheet with the background list of genes for these analyses is also included.

## Supplemental Data Legends

**Supplemental Data S1**. **Spotplots of domain clusters across samples.** Annotations are shown for each of 43 capture areas (generally 2 samples/tissue sections). Only spots retained after QC were assigned a domain; no annotation is plotted for spots removed by QC, revealing the H&E tissue beneath. Individual samples and annotations can be interactively viewed in uniform anatomic orientation using the provided resources in *Data Access and Visualization*.

**Supplemental Data S2**. **Spot deconvolution results using RCTD and previously described adult human LC-NE snRNA-seq**. An .*RDS* object is provided, with a named list object at the top level containing four different tabular results, where each row is a spot, each column is an snRNA-seq label, and each cell is an *RCTD*^92^ weight, giving the estimated proportion of spot expression attributable to an snRNA-seq cell type. The list names indicate the *RCTD* mode used (“Multi”--deconvolution assuming a spot contains ≤4 cell types from the reference data, or “Full”--deconvolution with no limit on the number of cell types) and snRNA-seq cell annotation used (“label”: 30 unnamed granular clusters; “label_merged”: 10 named, broad cell types grouped from these 30 clusters). For readability, the granular ‘labels’ were concatenated with the cell type they were grouped with in ‘label_merged’ (e.g., the granular label 4 was grouped into ‘inhibitory’ under label_merged, and thus is presented here as “inhibitory_4”).

**Supplemental Data S3**. **Complete *SCORPION* GRN for LC^NM+^ and LC^NM-^ with edge weight comparisons.** Unabridged version of the GRN edge comparisons in **Table S8**. Columns *nmp* and *nmn* provide the *Z*-normalized edge weights from LC^NM+^ and LC^NM-^ GRNs, respectively. Column *diff* is the edge weight difference, and nmp_spec/nmn_spec indicate edges determined to be specific to LC^NM+^ or LC^NM-^ (see *Methods*), respectively.

**Supplemental Data S4**. **Raw NMF matrices *w* and *h* resulting from decomposition of the LC domain into 160 factors.** An .*RDS* object is provided containing a list with two slots, “*w*” and “*h*”, corresponding to gene weight and factor loading matrices. Factor, gene, and spot identifiers are included as row/column names.

**Supplemental Data S5**. **Complete *Enrichr***^105^ **results from analysis of genes associated with NM intensity or NM pixel proportion.** An .RDS list object with named slots containing the concatenated results from NM intensity or NM pixel proportion tests (6 tests per NM metric: FDR-significant or Bonferroni-significant genes and subsets thereof in each direction of association). The same tables filtered to top enrichments and with a key with more information for interpretation are in **Table S14**.

