## Supplemental Figures S1-S19 for "Impact of Alzheimer’s disease risk factors and local neuromelanin content on the transcriptomic landscape of the human locus coeruleus"

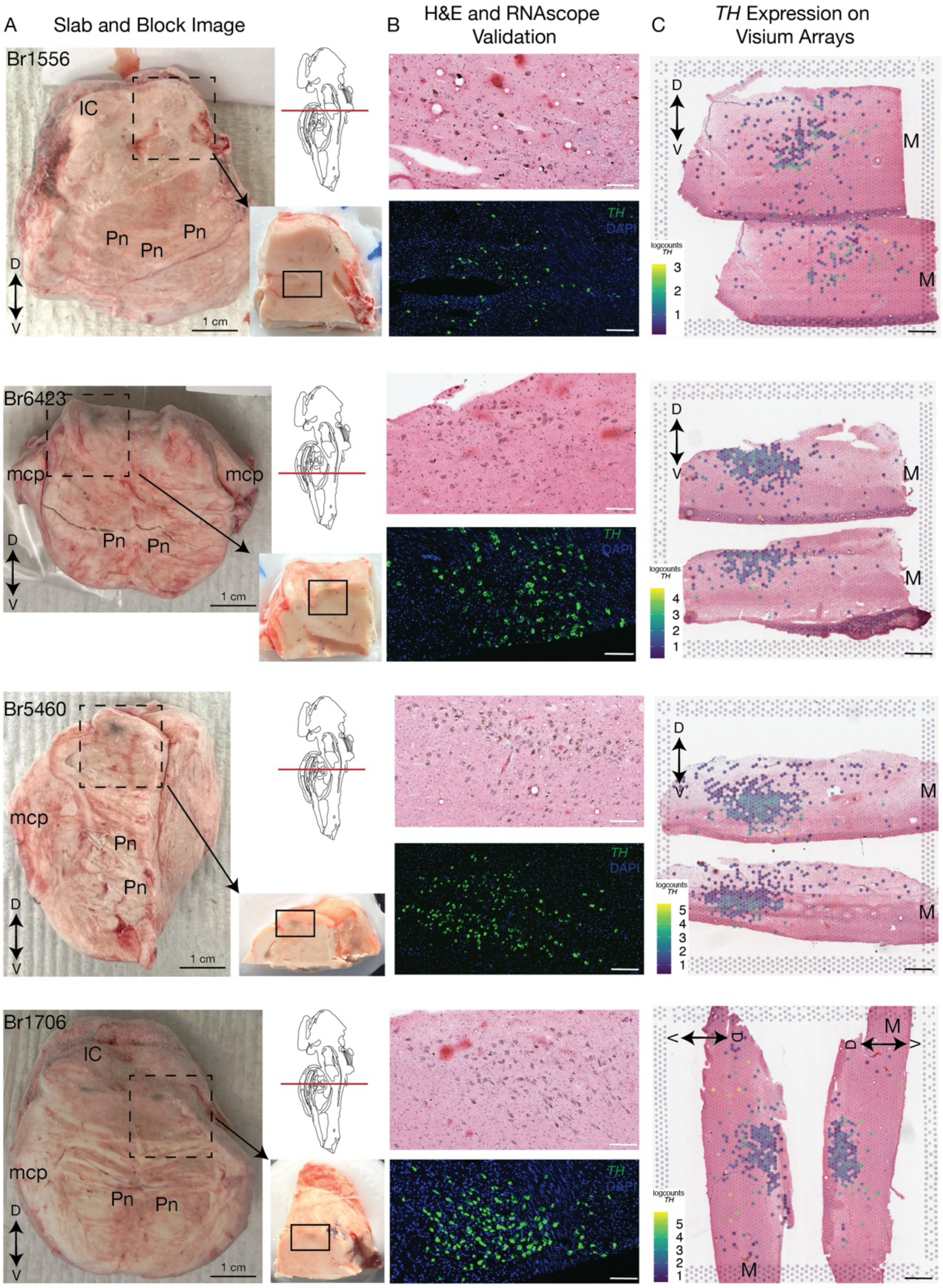


**Supplemental Figure S1. Neuroanatomical validation of LC inclusion and orientation on tissue blocks.** **A)** Images of fresh-frozen transverse slabs of postmortem human brain at the level of the dorsal pons from n=4 representative donors. Images are annotated with surrounding neuroanatomical features including the middle cerebellar peduncle (mcp), Inferior Colliculus (IC) and the pontine nucleus (Pn). Dashed boxes indicate the location of the tissue block that was dissected from the slab, with arrows pointing to the image of the tissue block. The solid black box indicates the boundaries of scored tissue placed on the Visium capture area in **C**. Above the image of the tissue block is a brain atlas schematic corresponding to the level of the transverse cut of the brain slab. **B)** Summary of anatomical validation experiments for respective tissue blocks shown in **A**. Brightfield images of H&E staining (upper) and fluorescent images from single molecule fluorescent *in situ* hybridization (smFISH, lower) with DAPI (blue) and *TH* (green). **C)** Sections from the tissue blocks in **A** after mounting to Visium capture areas and H&E staining. Log counts of *TH* gene expression from SRT analysis are overlaid on tissue sections. Neuroanatomical orientation is indicated by arrows: D - dorsal; V - ventral; M - medial.

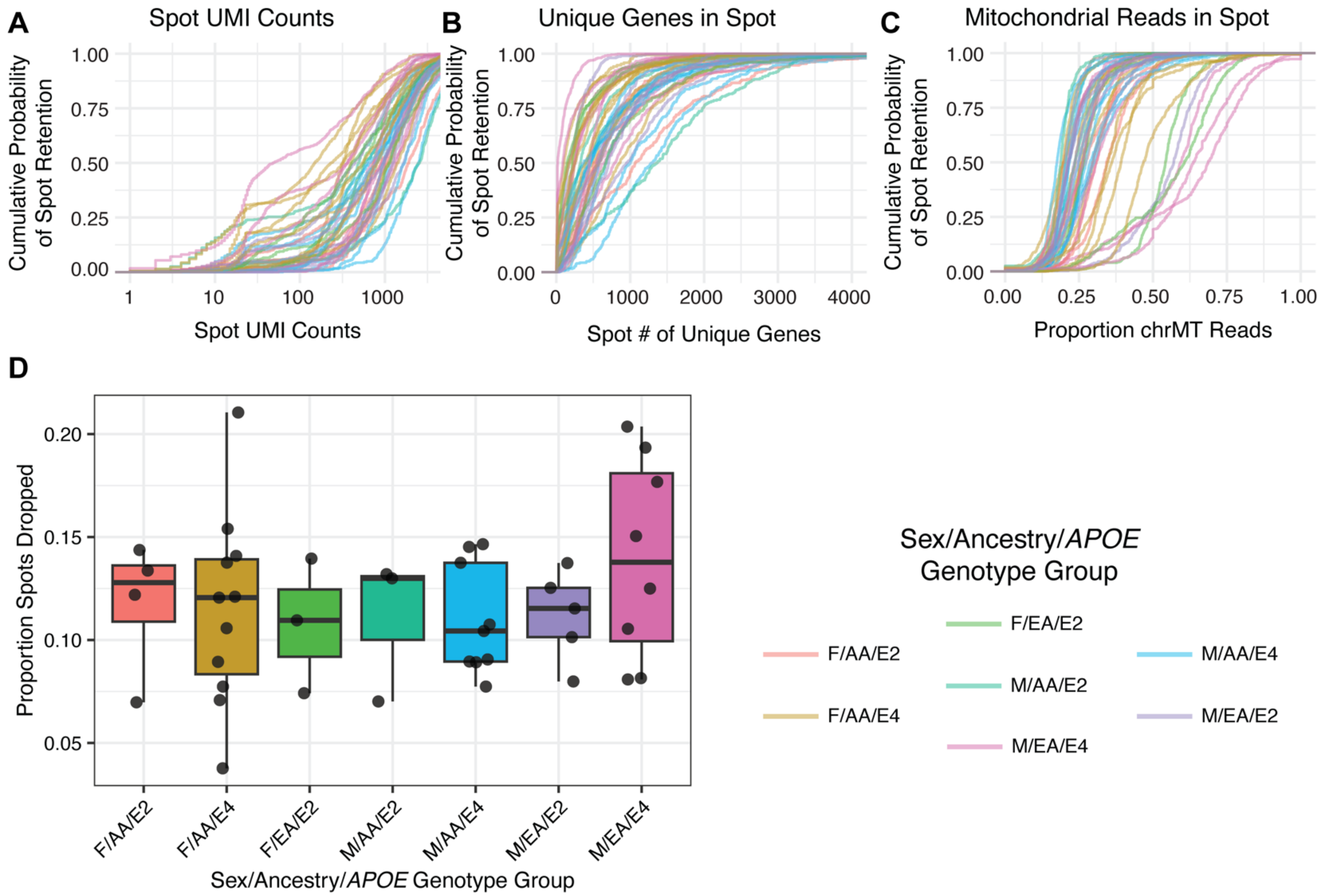


**Supplemental Figure S2. Spots removed by QC steps prior to analysis.** Cumulative probabilities of tissue-overlapping spots passing QC are illustrated as lines, one line per capture areas (43 total) and colored by ancestry-sex-*APOE* genotype group, for 3 standard QC metrics: **A)** UMI count; **B)** number of unique genes in the spot; and **C)** proportion of mitochondrial reads in the spot. **D)** The proportion of within-tissue spots discarded during QC are shown per capture area for each ancestry-sex-*APOE* genotype group. (This includes the outermost perimeter of spots classified as tissue-overlapping, which were removed indiscriminately during the QC process). Legend applies to all panels.

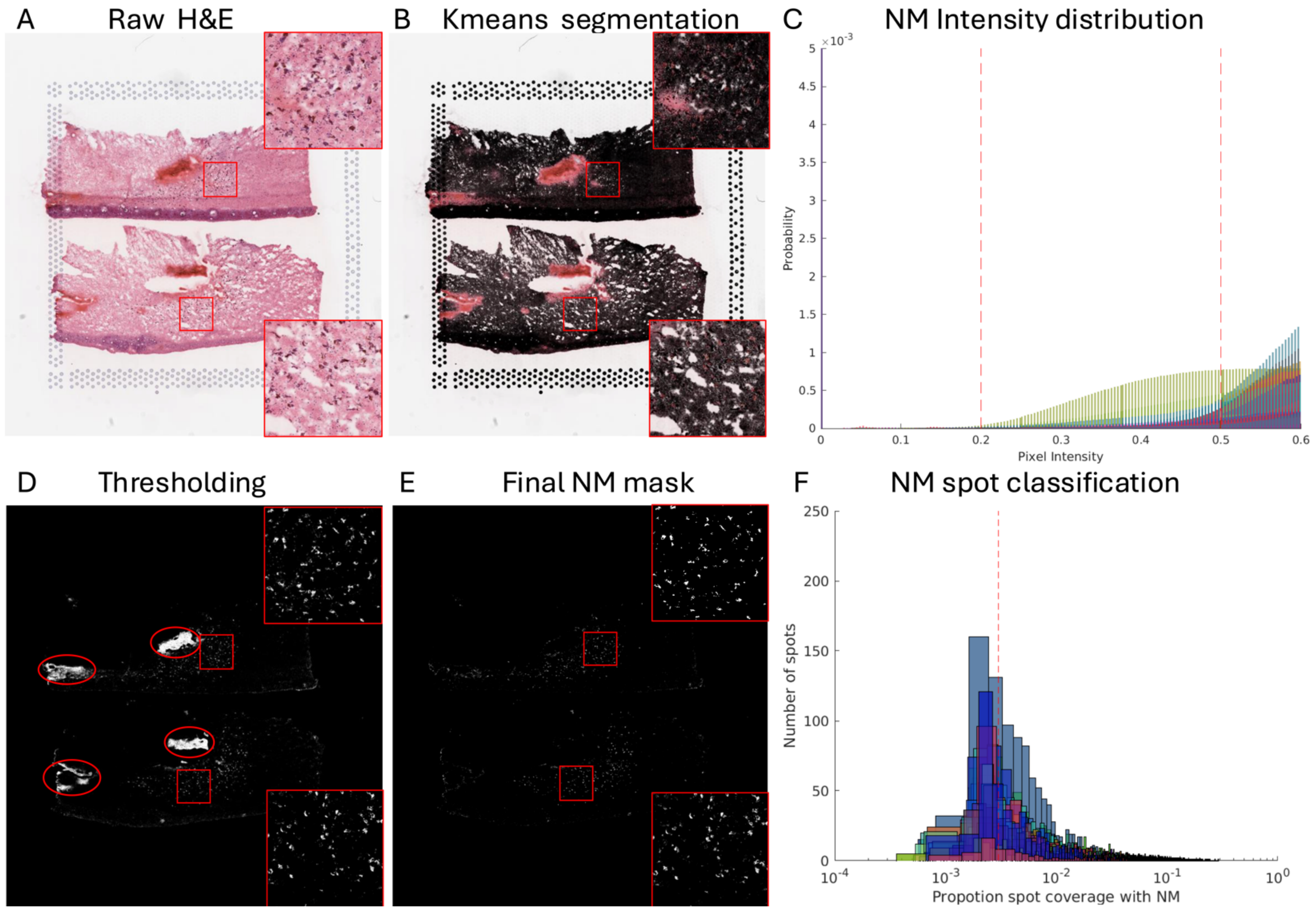


**Supplemental Figure S3. Neuromelanin (NM) segmentation workflow. A)** High-resolution raw H&E image with insets showing NM pigment in LC region. **B)** *k*-means segmentation of **A** using *VistoSeg::VNS* function. **C)** Intensity distribution of Kmeans segmented regions from **B** for all samples. **D)** Thresholding applied to the grayscale-converted image from **B** to extract regions with intensities between 0.2 and 0.5, resulting in the detection of both tissue artifacts (highlighted in ovals) and NM regions whose colors overlap. **E)** Final NM segmentation obtained after post-processing to remove tissue artifacts identified in **D**. **F)** Distribution of the proportion of NM pixels per Visium spot across all samples, with a 0.3% threshold to classify spots as NM+ (above threshold) or NM- (below threshold).

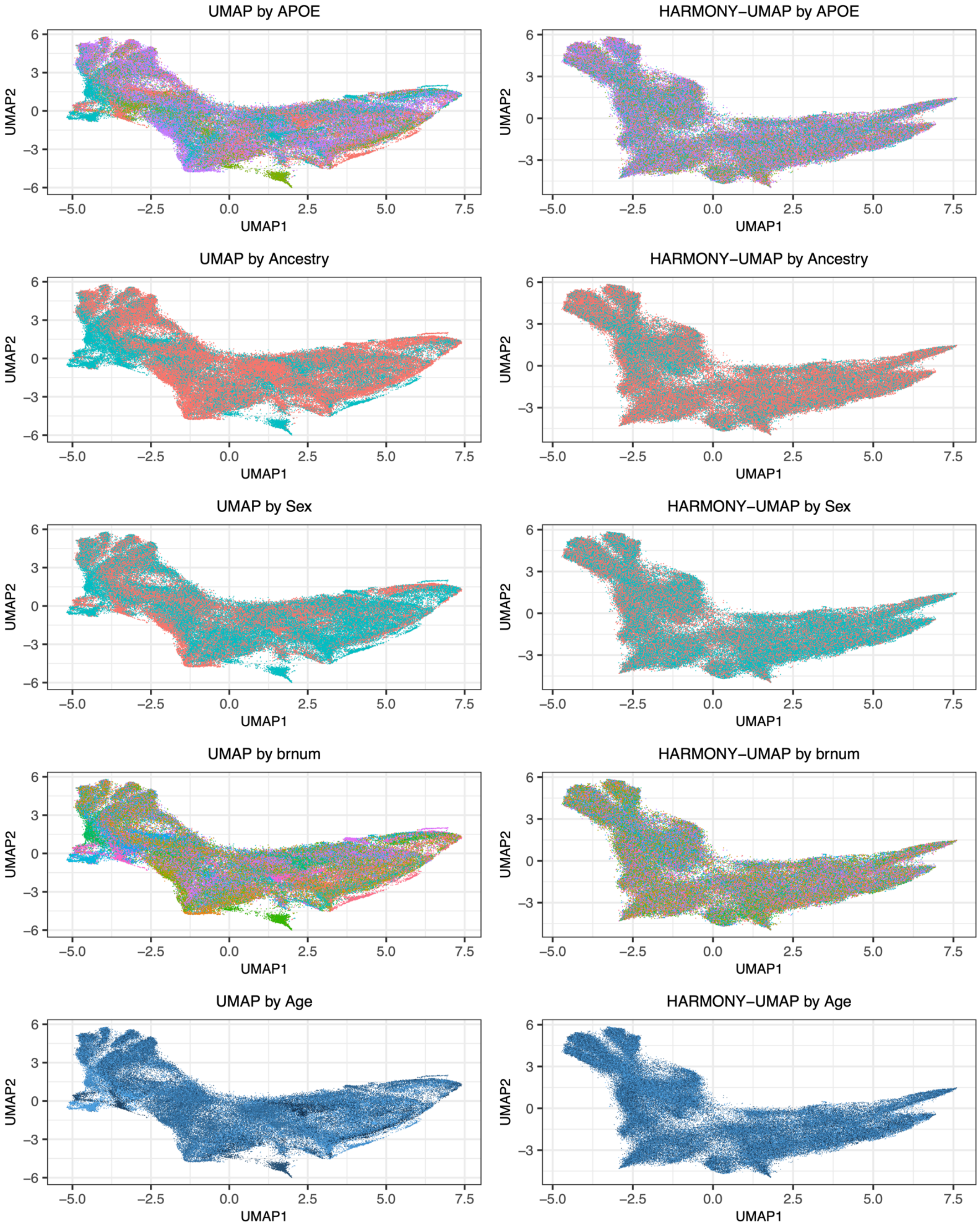


**Supplemental Figure S4. UMAP before and after *Harmony* integration, coded by donor or donor characteristics**. Each plot shows a UMAP of the SRT data from 33 donors before batch correction with *Harmony*[77]. Plots are titled by the variable they illustrate, with points colored by group for categorical variables, or dark blue to light for continuous variables. For a given variable, the initial UMAP is color-coded and displayed on the left, and UMAP after *Harmony* integration coded by the same variable is shown to its right.

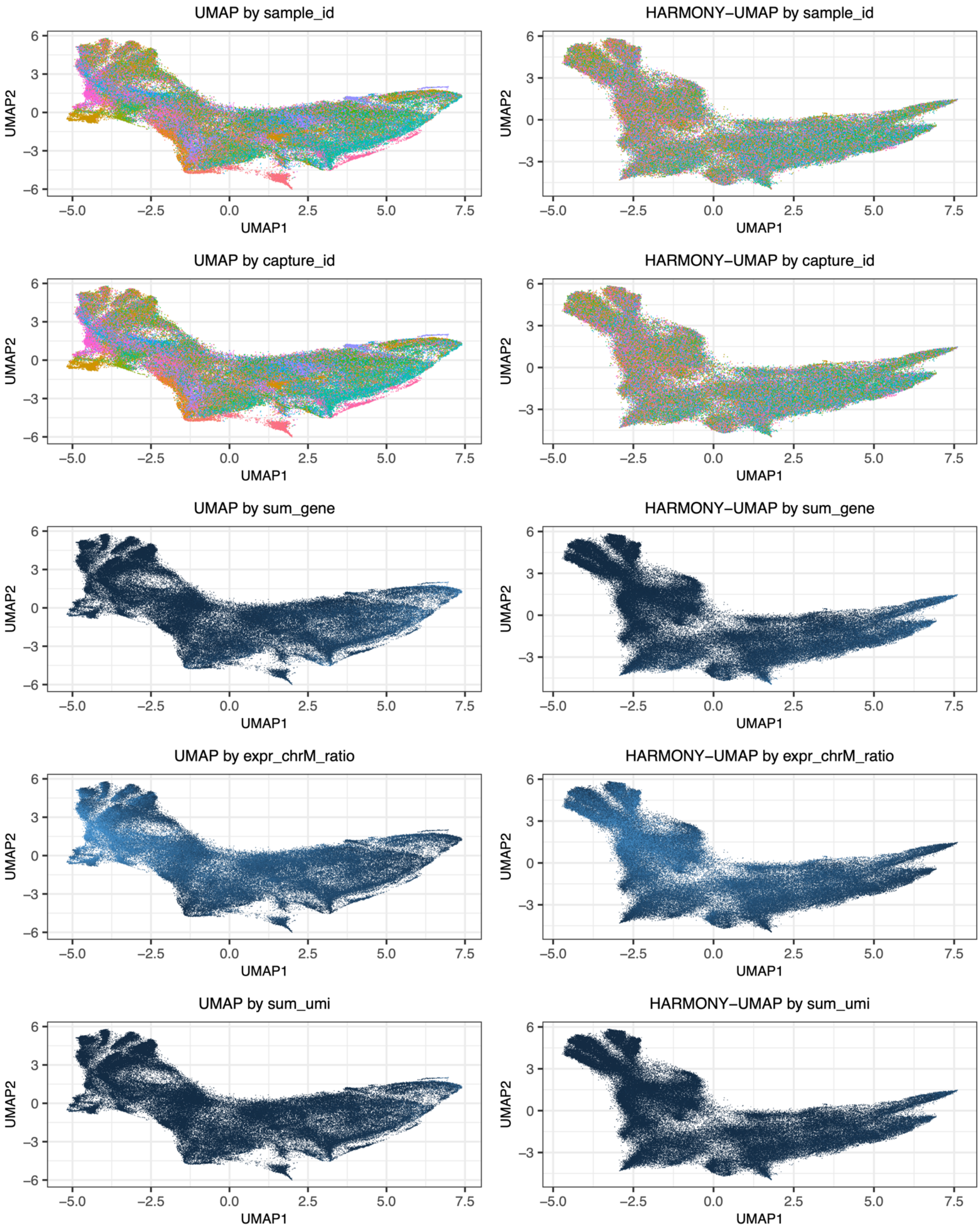


**Supplemental Figure S5. UMAP before and after *Harmony* integration, coded by technical variables**. Each plot shows a UMAP of the SRT data from 33 donors before batch correction with *Harmony*[77]. Plots are titled by the variable they illustrate, with points colored by group for categorical variables, or dark blue to light for continuous variables. For a given variable, the initial UMAP is color-coded and displayed on the left, and UMAP after *Harmony* integration coded by the same variable is shown to its right.

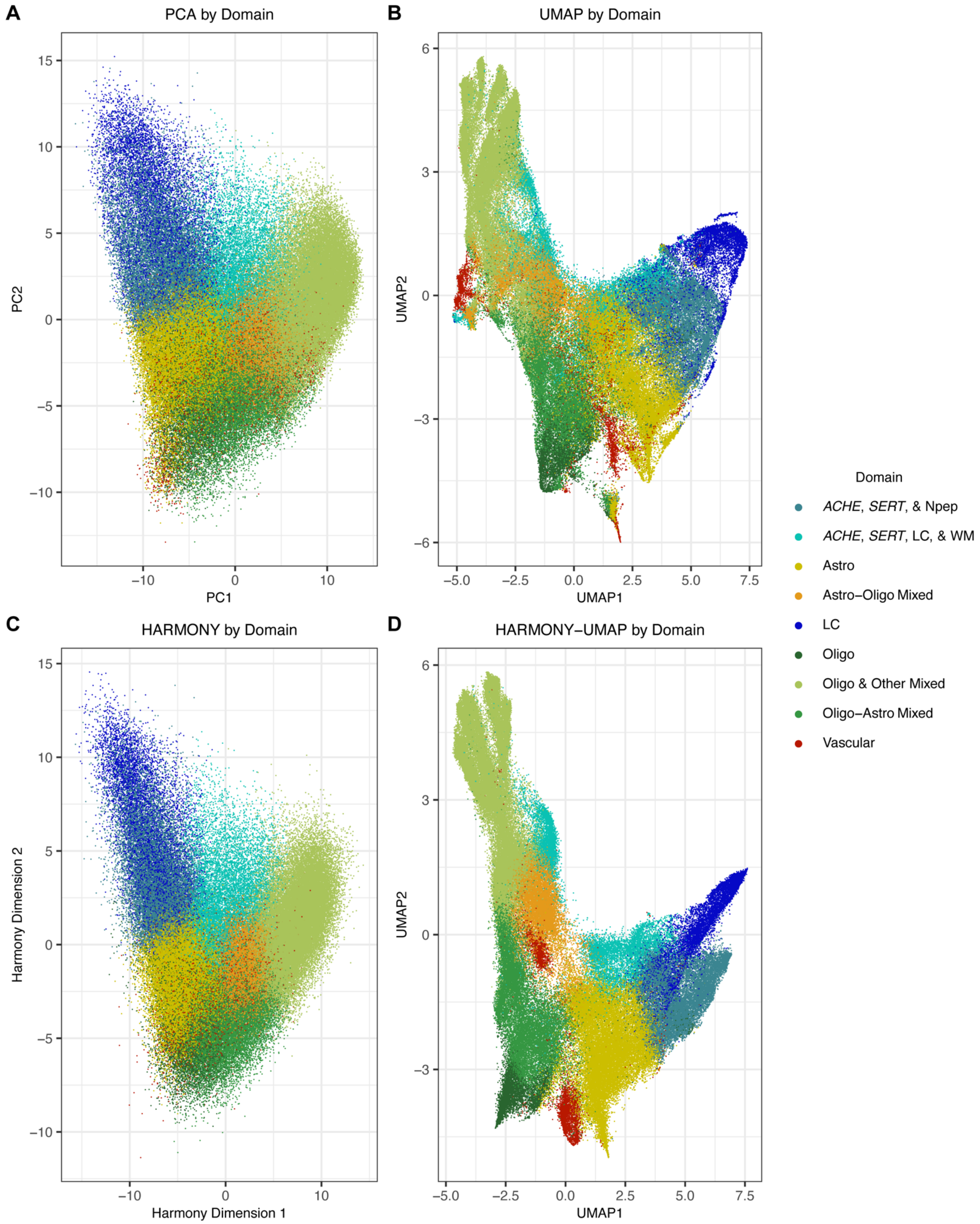


**Supplemental Figure S6. Dimensionality reductions before and after batch correction, coded by domain. A)** Uncorrected PCA plot. **B)** UMAP of the uncorrected PCA. **C)** *Harmony*[77] batch-corrected dimensionality reduction, dimensions 1 and 2. **D**) UMAP of the full *Harmony*-corrected dimensionality reduction.

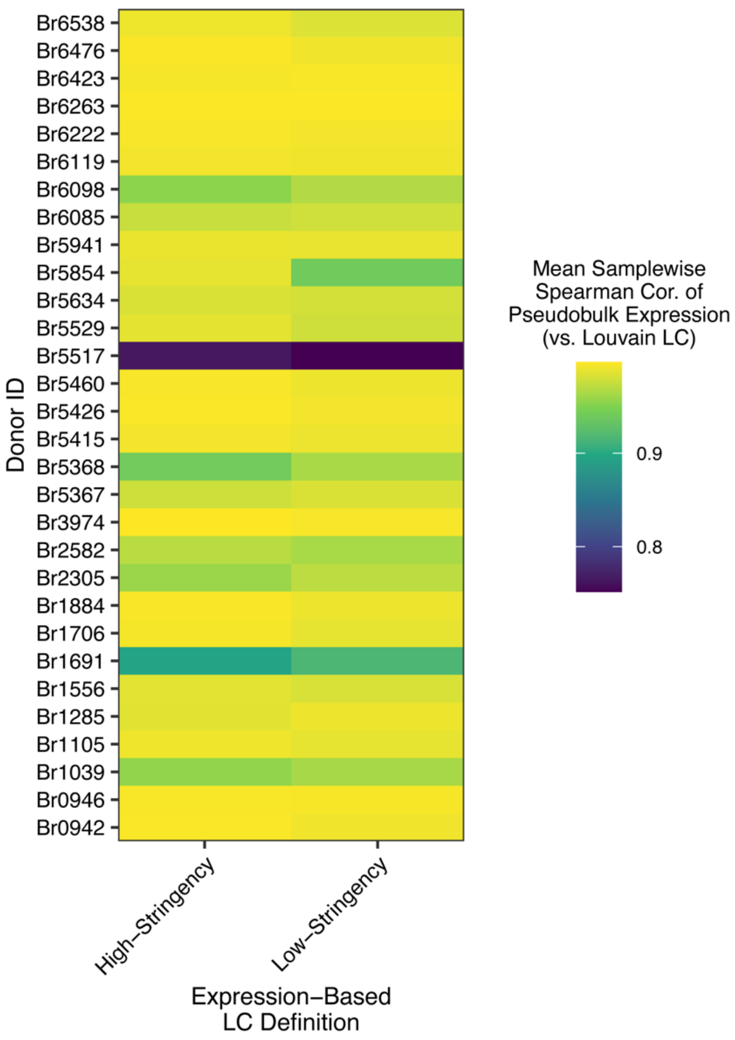


**Supplemental Figure S7. Donor-wise agreement between Louvain-clustered LC domain and expression-based definitions of LC spots.** For each sample, LC spots were manually labeled based on LC marker genes. The low-stringency condition required ≥1 count of *DBH*, ≥1 count of either *SLC6A2* or *TH*, and ≥1 count of additional LC marker genes (*PHOX2A*, *PHOX2B*, *SLC18A2*, *DDC*). The high-stringency definition required that the spot have ≥5 counts combined of *DBH*/*TH*/*SLC6A2 and* ≥2 counts of additional markers. LC as defined manually or by Louvain clustering was then pseudobulked for each donor, and the Spearman correlation of logcounts in the manual pseudobulks calculated relative to the LC domain.

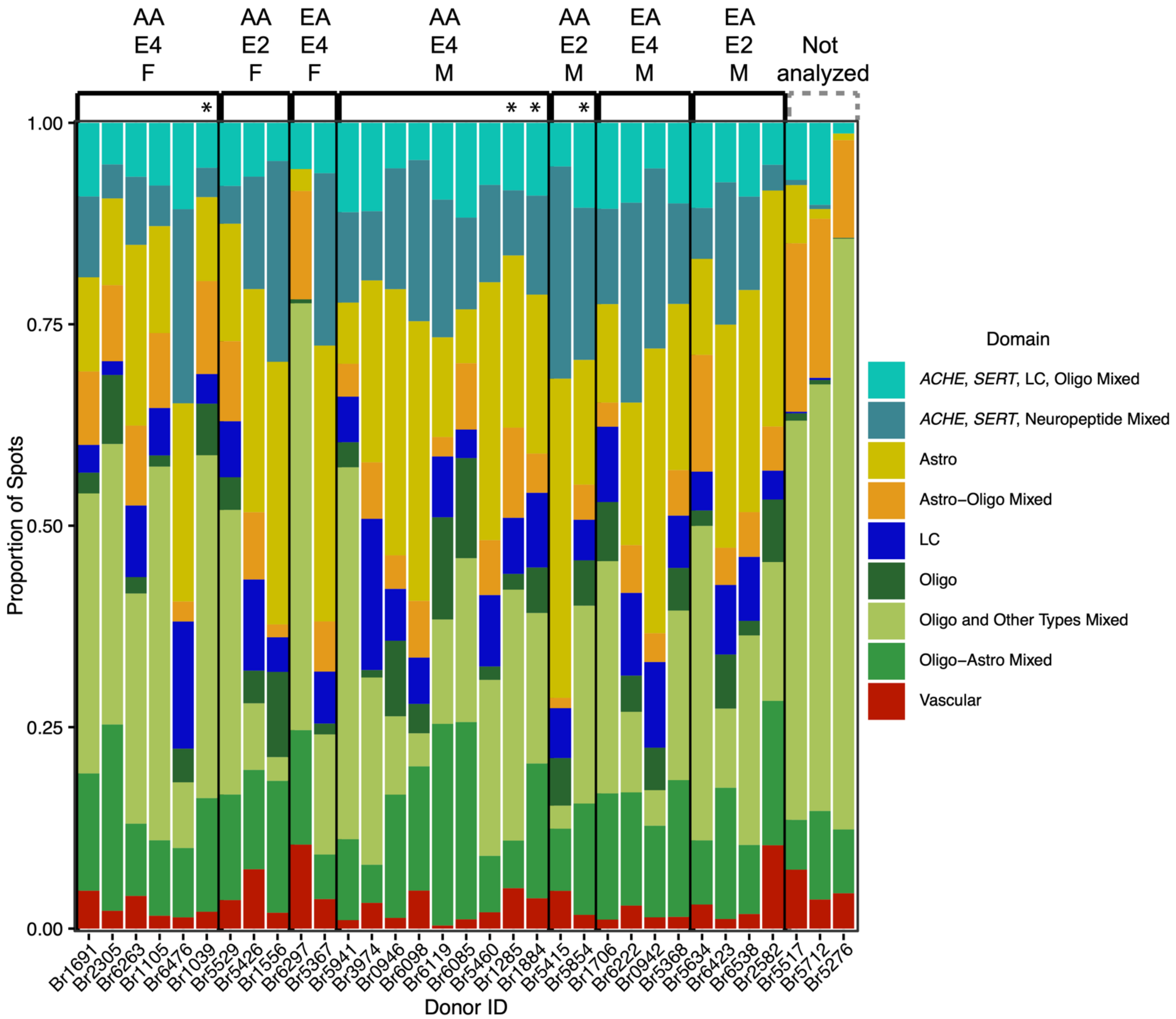


**Supplemental Figure S8. Proportion of spots in each domain by donor**. Stacked barplot showing the proportion of analyzed SRT spots from one donor (*x*-axis) assigned to a given domain. Bars are grouped by ancestry, *APOE* genotype, and sex group. Donors for which SRT samples were determined to be outside of LC territory are shown at far right. * denotes that this donor was excluded from categorical ancestry-genotype DE analyses due to AA genomic ancestry proportion below <75%.

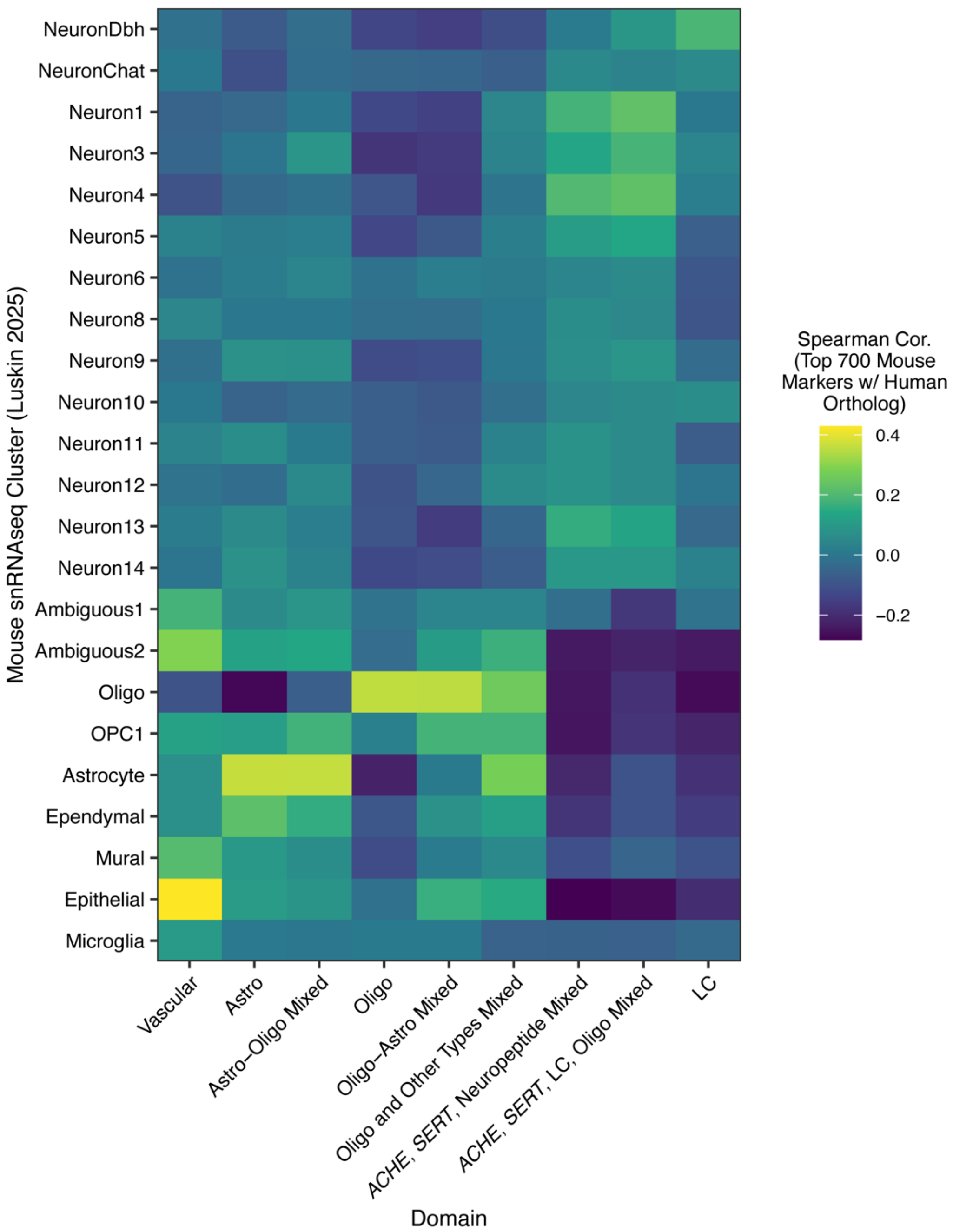


**Supplemental Figure S9. Spatial registration[87-89] of human LC domains to mouse snRNA-seq of the LC and peri-LC.** While most SRT domains showed the strongest registration to the expected mouse cell type, *ACHE-SERT* domains could not be assigned to any one mouse peri-LC neuron cluster. Heatmap showing cluster registration between SRT domains and 25 mouse snRNA-seq clusters reported in[86]. Only genes represented in both datasets with 1-1 orthologs were used. Registration calculations used the top 700 markers for each mouse cluster (the same calculations using a range of 10 to 5000 top-enriched genes per mouse cluster are in **Table S2**).

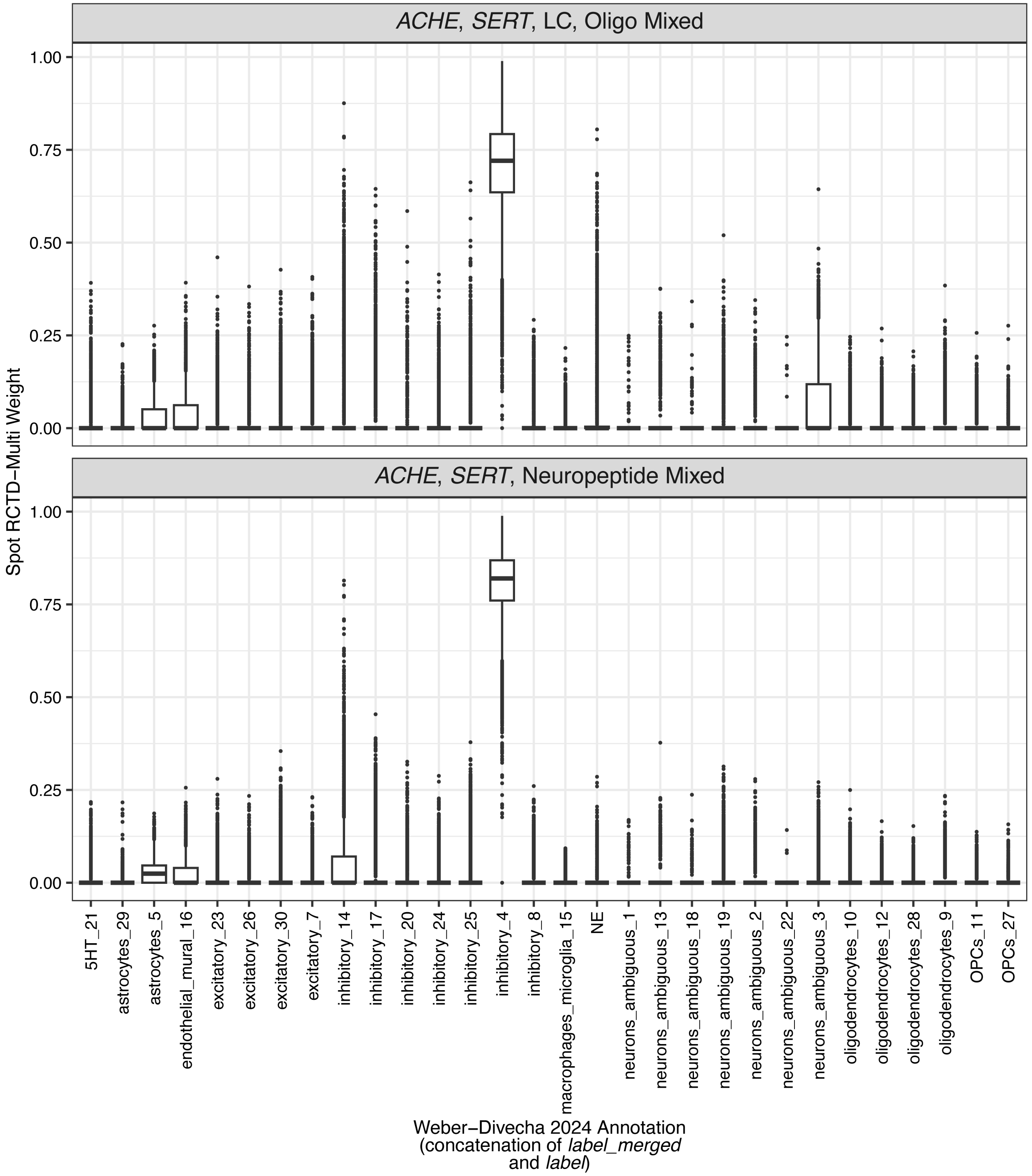


**Supplemental Figure S10. Spot deconvolution suggests *ACHE-SERT*-coexpressing domains are rich in one inhibitory cell type identified in human LC snRNA-seq**. Spot deconvolution of SRT data using snRNA-seq profiles from human LC[90] was performed using the 29 initial clusters defined in the snRNA-seq data. Plotted are the spot-level *RCTD* weights (estimated cell type proportions) for spots in each *ACHE-SERT* domain. The "inhibitory 4" snRNA-seq cell type is estimated to underlie 50-75% of transcriptomic signal in nearly all SRT spots from both *ACHE-SERT* domains. Marker analysis was then performed for 'inhibitory 4' relative to all other inhibitory clusters in the snRNA-seq data (**Table S5**). Full results of *RCTD*[91] analysis using these 29 snRNA-seq clusters (and using the 10 broader types, 'label_merged', that the authors grouped these into) are in **Data S2**.

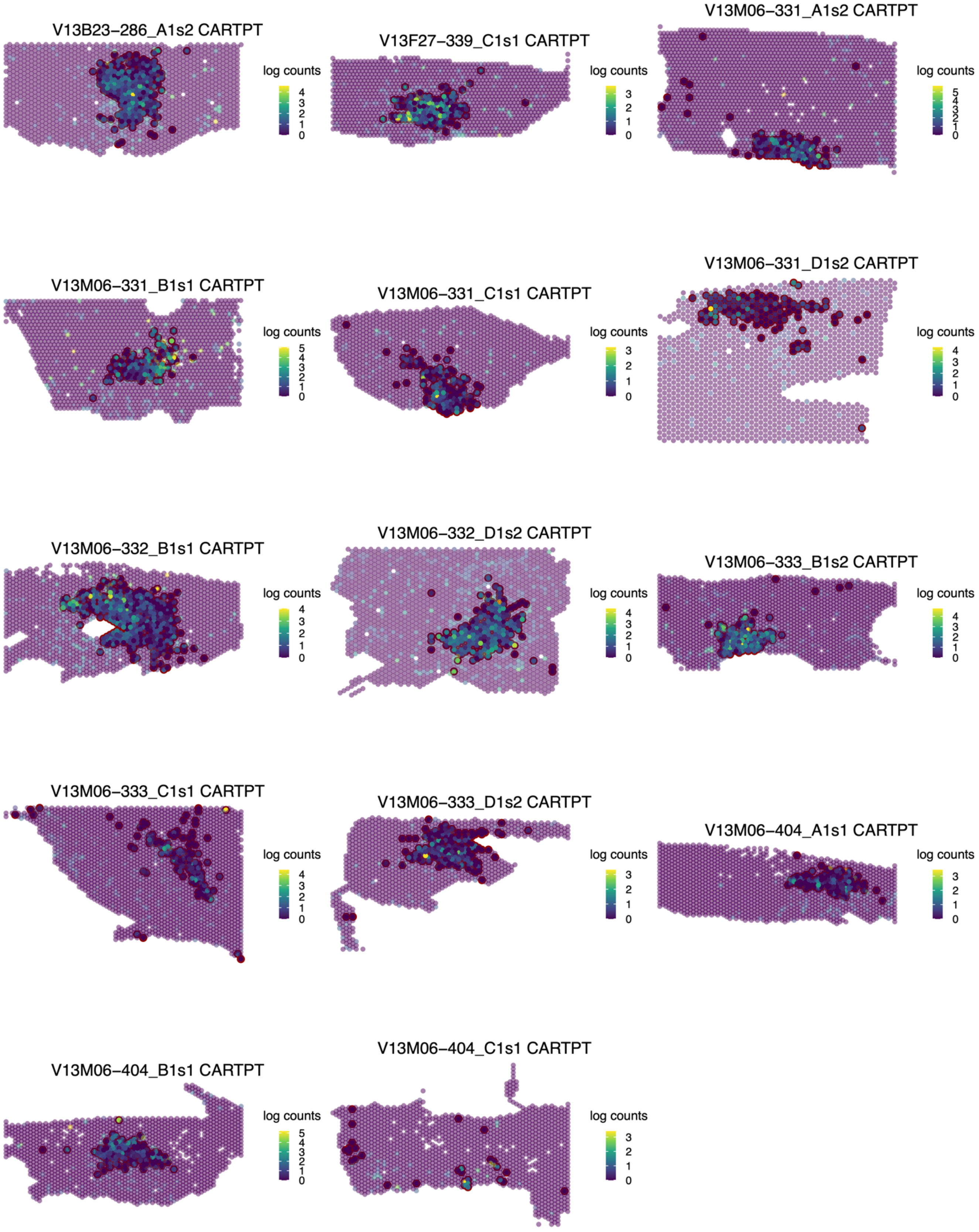


**Supplemental Figure S11. Spatially variable *CARTPT* expression along the medial-lateral axis of the LC.** *CARTPT* expression is plotted for samples for which the gene was significantly spatially variable (uncorrected $p<0.05$; only one sample plotted per donor). All samples are plotted with their medial edges at left and dorsal edges at bottom.

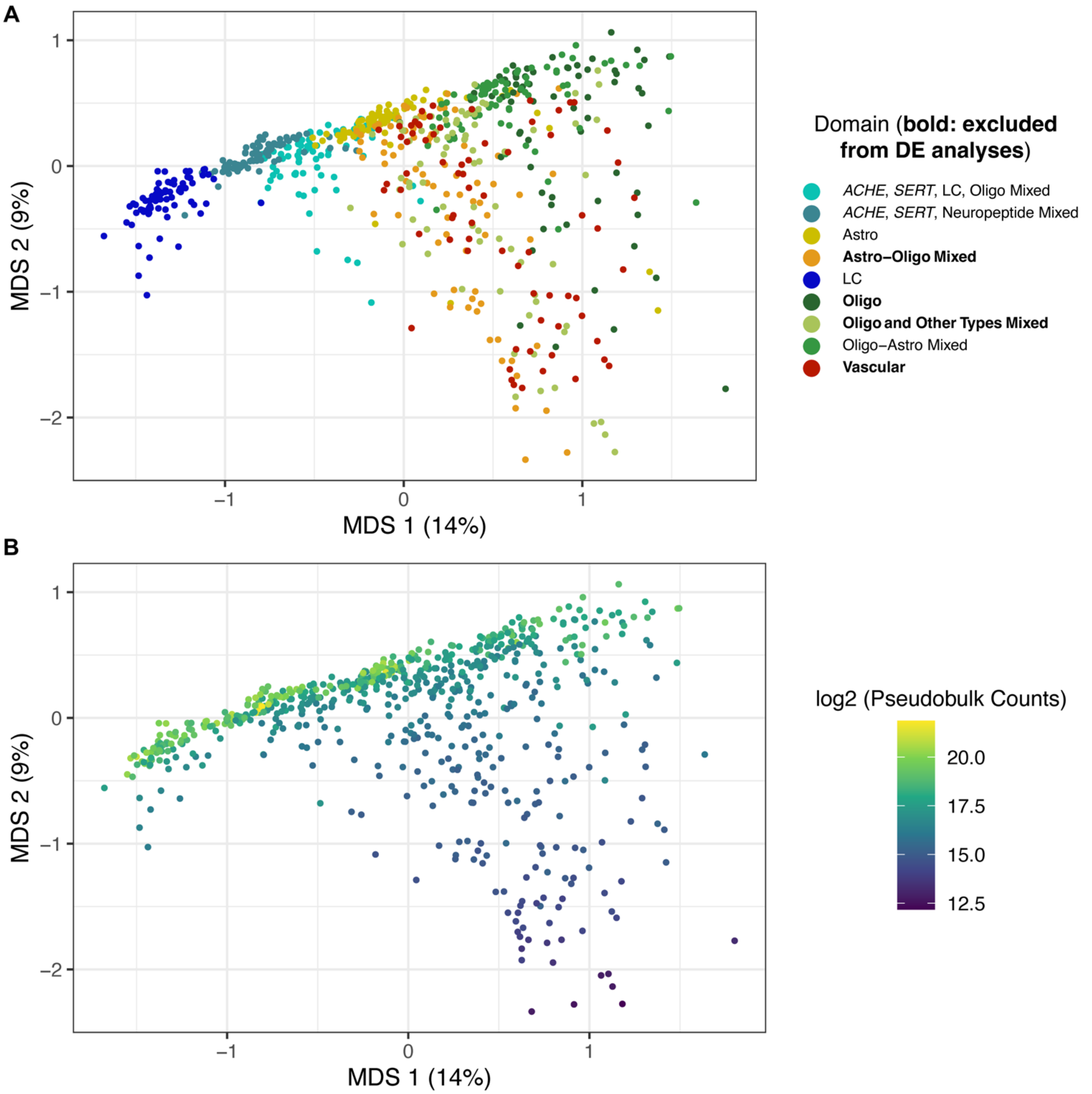


**Supplemental Figure S12. Multidimensional scaling (MDS) plots of pseudobulked data for DE analyses**. **A)** Each sample (one domain from one tissue section) in MDS space, color-coded by domain, notable for a second dimension separating the domains excluded from DE analyses (Oligo, Astro-Oligo, Vascular, and Oligo-Other Cell Types). **B)** The same plot, color-coded by read count in the pseudobulk sample.

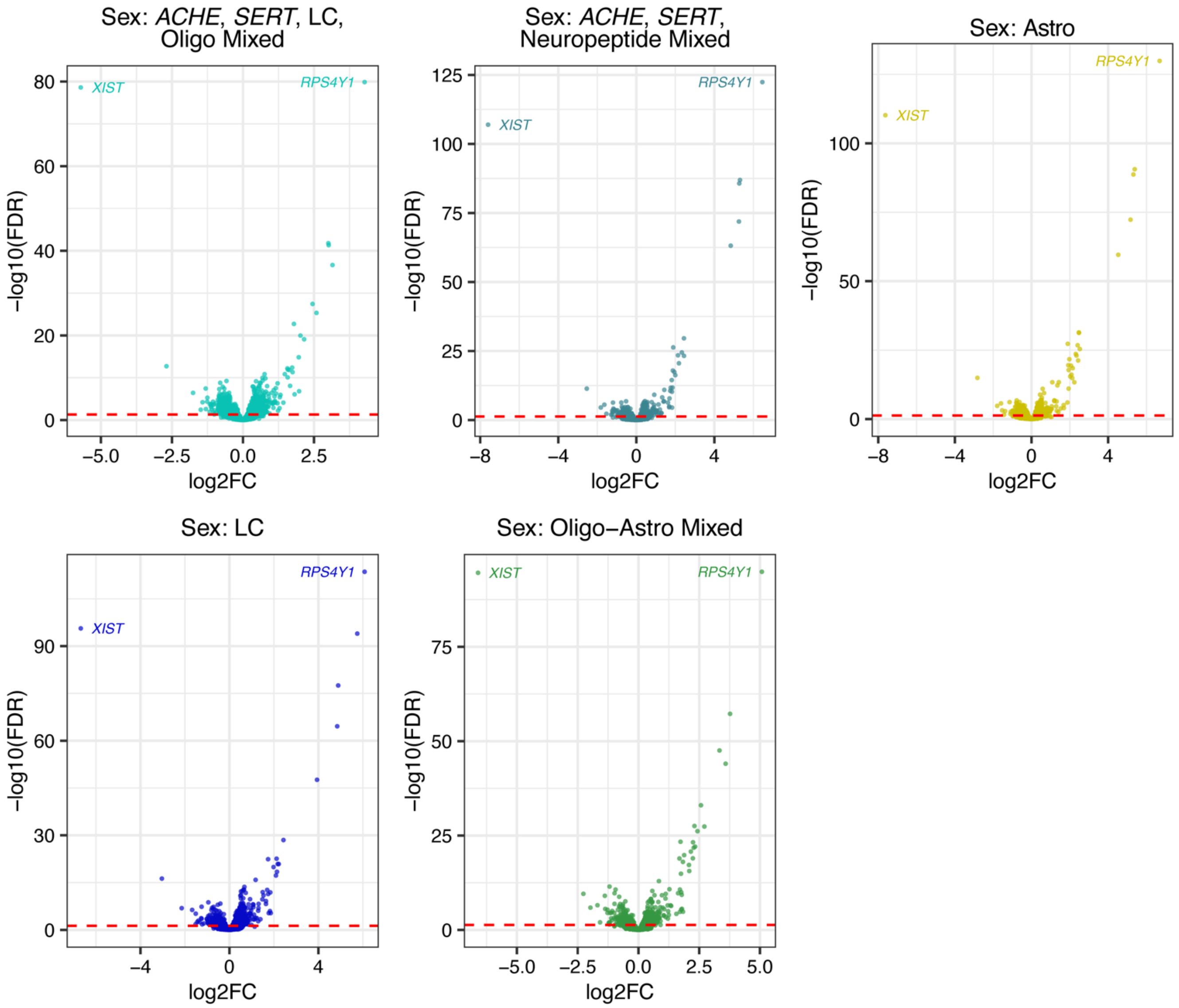


**Supplemental Figure S13. Sex-DE volcano plots from each domain analyzed.** Sex DE volcano plots, including sex chromosomal genes, are shown. The top DEGs are labeled. Positive $\log_2$ fold change (logFC) signifies higher expression in males compared to females.

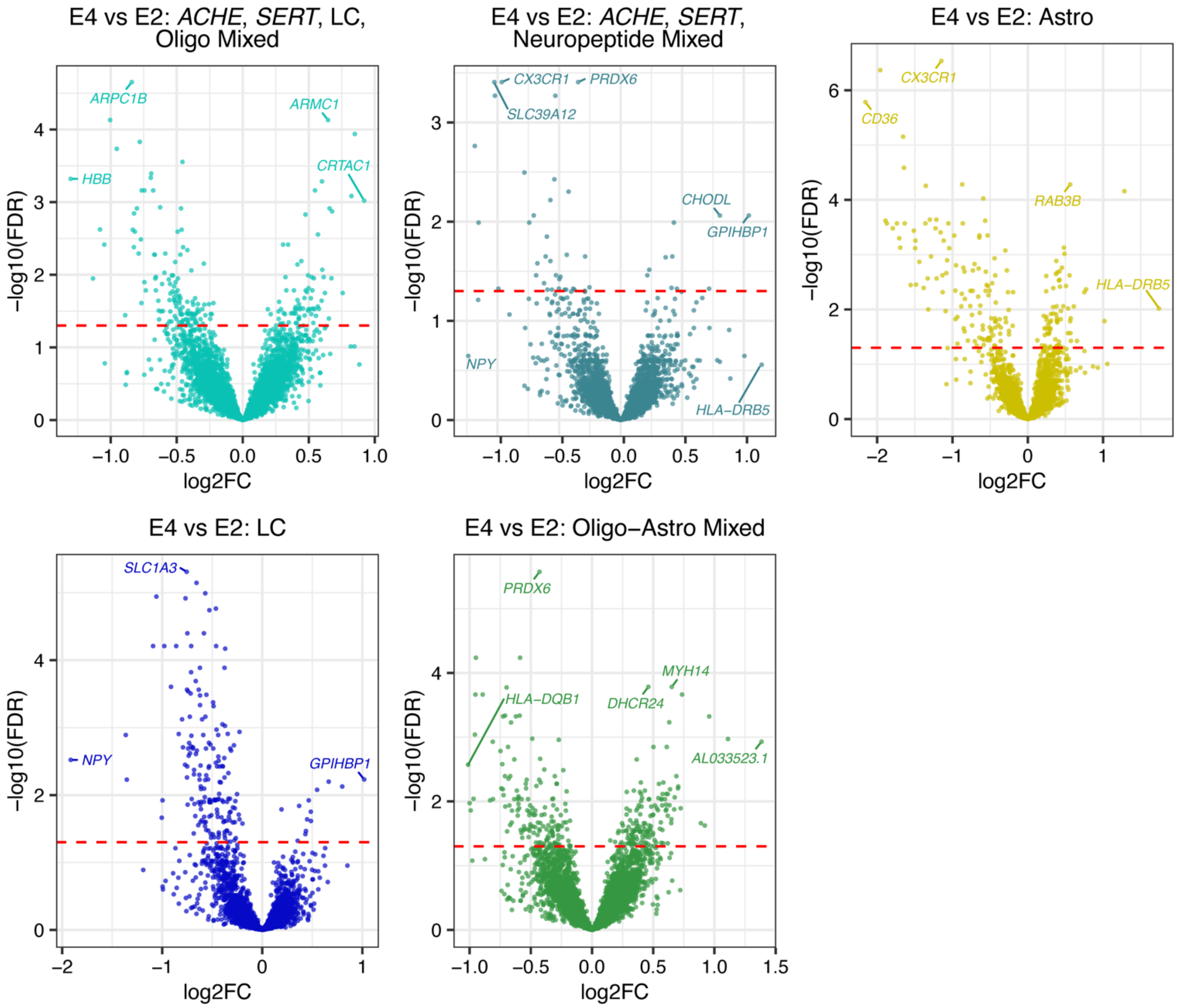


**Supplemental Figure S14. Genotype-DE volcano plots for each domain analyzed.** DE analysis between all E4 carriers and all E2 carriers. Top DEGs are labeled. Positive logFC signifies higher expression in E4 carriers compared to E2 carriers.

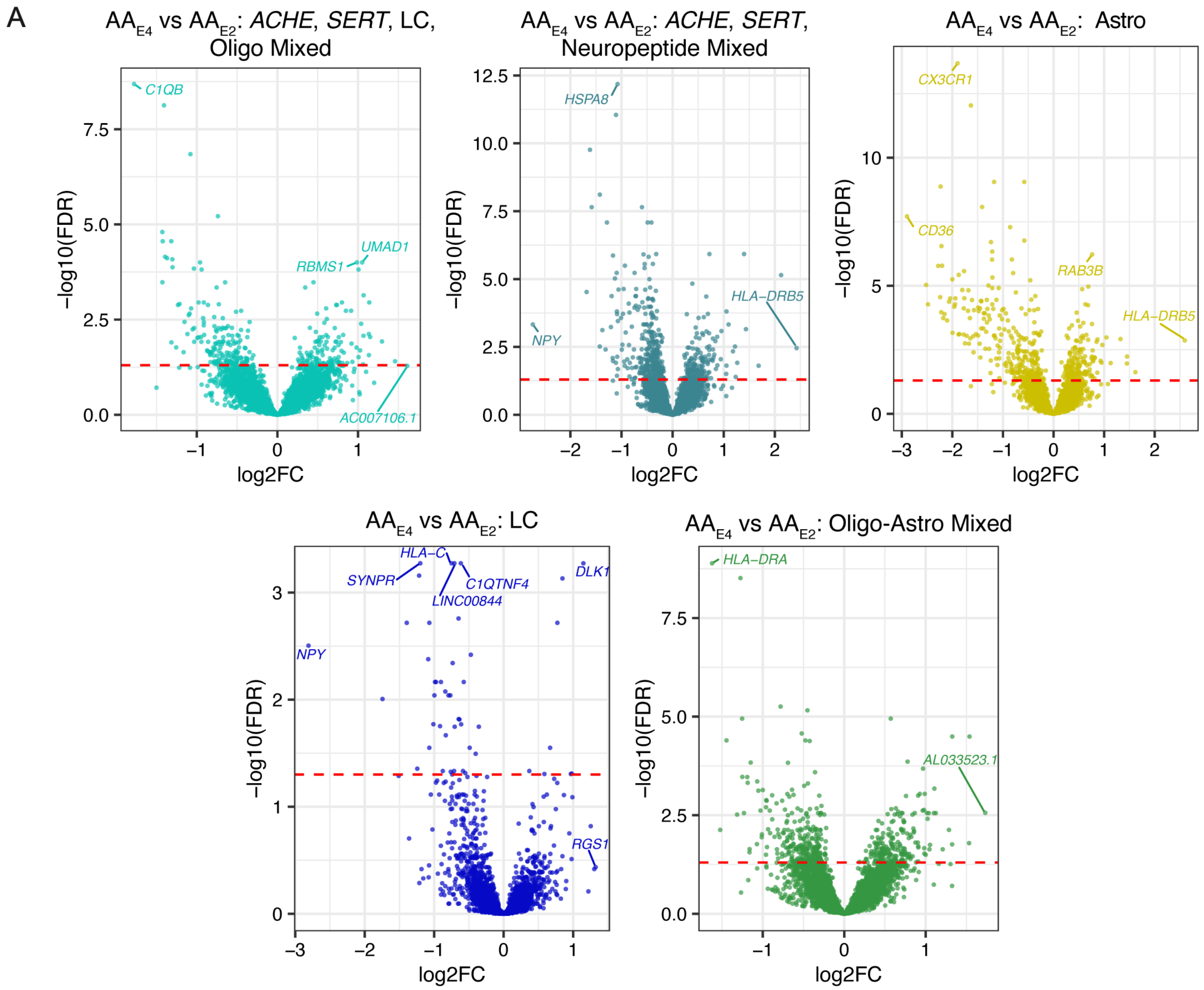


**Supplemental Figure S15. DE between E4 and E2 carriers of African genomic ancestry.** Top DEGs are labeled. Positive logFC signifies higher expression in E4 carriers compared to E2 carriers.

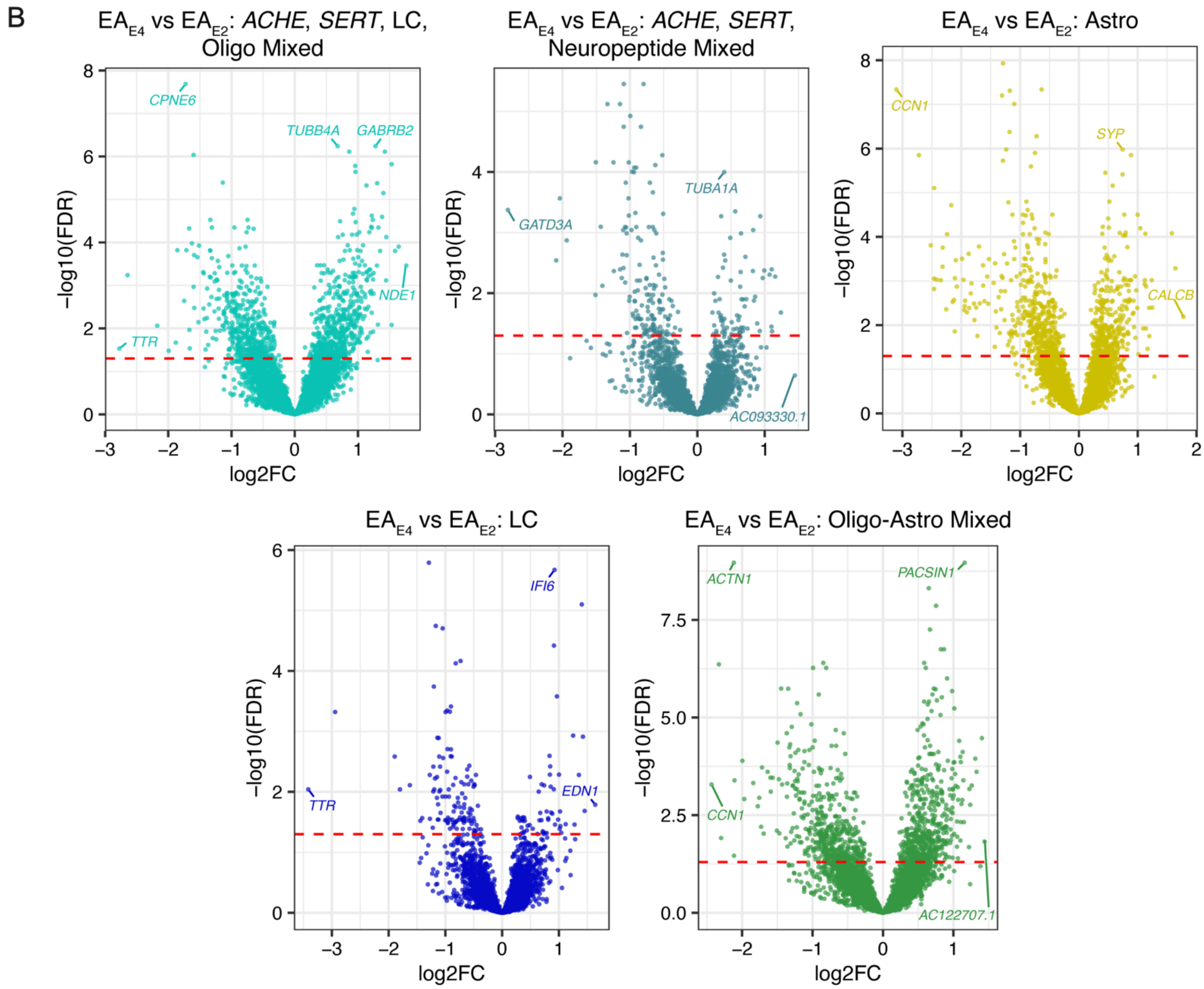


**Supplemental Figure S16. DE between E4 and E2 carriers of European genomic ancestry.** Top DEGs are labeled. Positive logFC signifies higher expression in E4 carriers compared to E2 carriers.

C

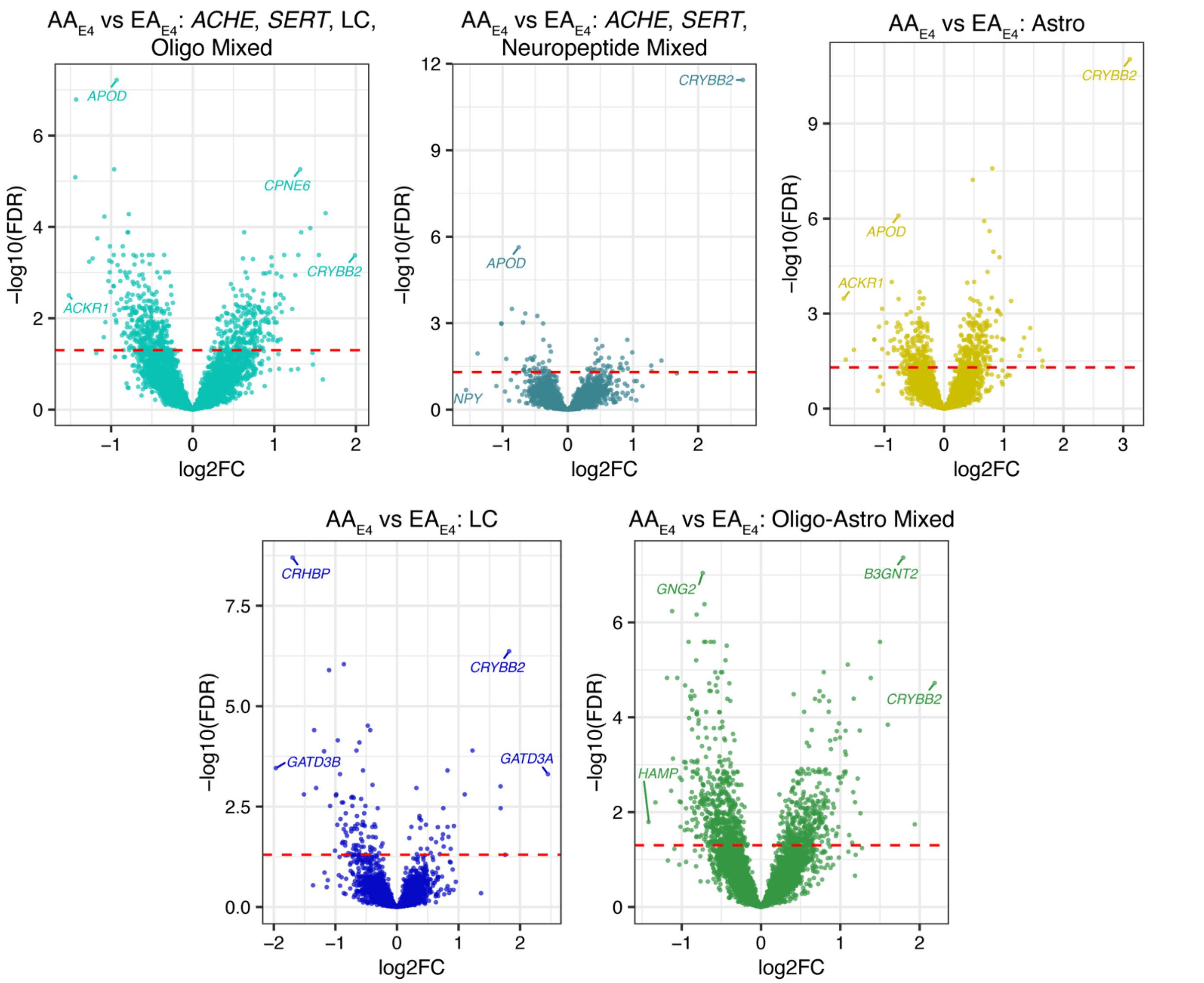


**Supplemental Figure S17. DE between African ancestry E4 carriers and European ancestry E4 carriers.** Top DEGs are labeled. Positive logFC signifies higher expression in AA E4 carriers compared to EA E4 carriers.

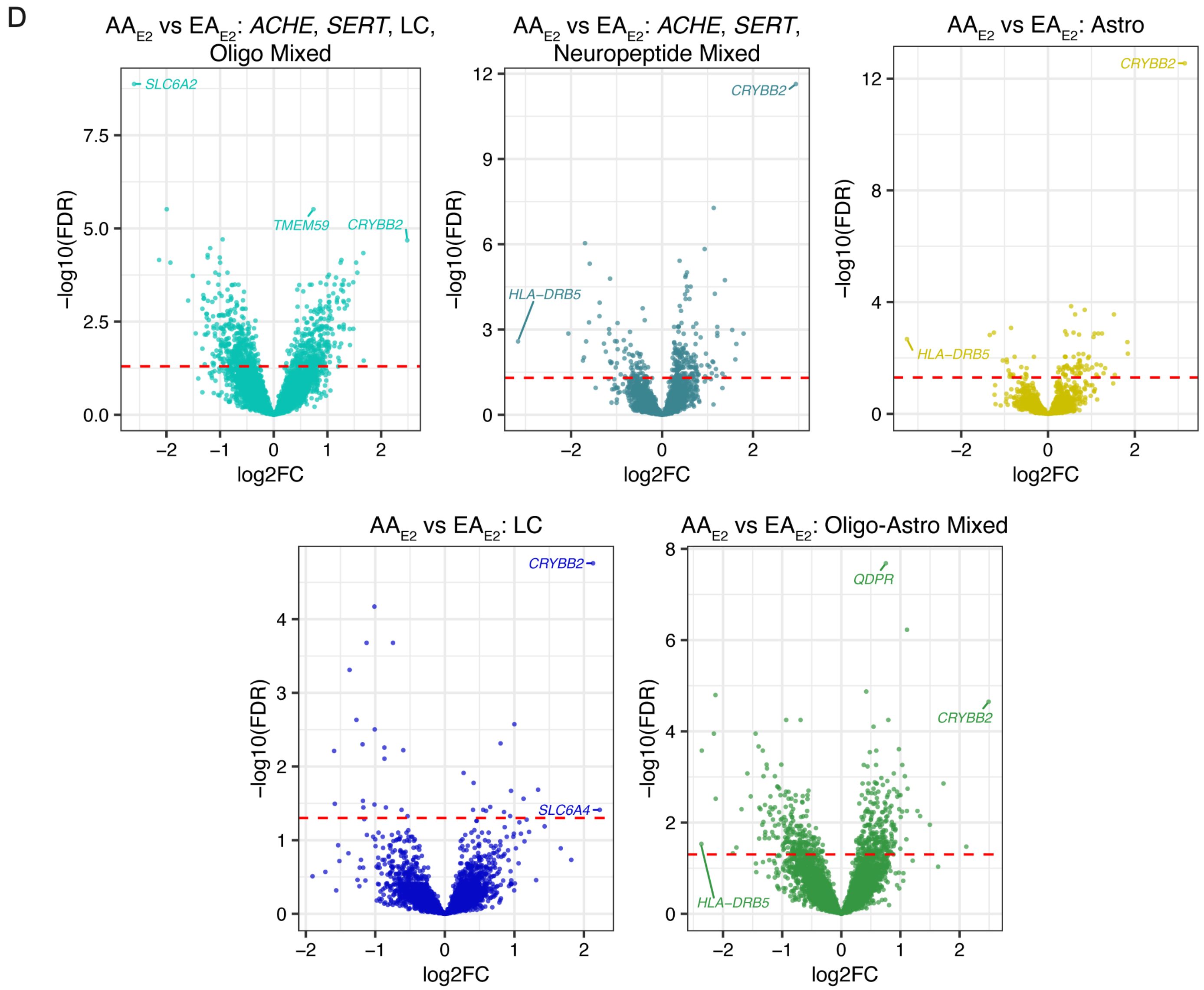


**Supplemental Figure S18. DE between African ancestry E2 carriers and European ancestry E2 carriers.** Top DEGs are labeled. Positive logFC signifies higher expression in AA E2 carriers compared to EA E2 carriers.

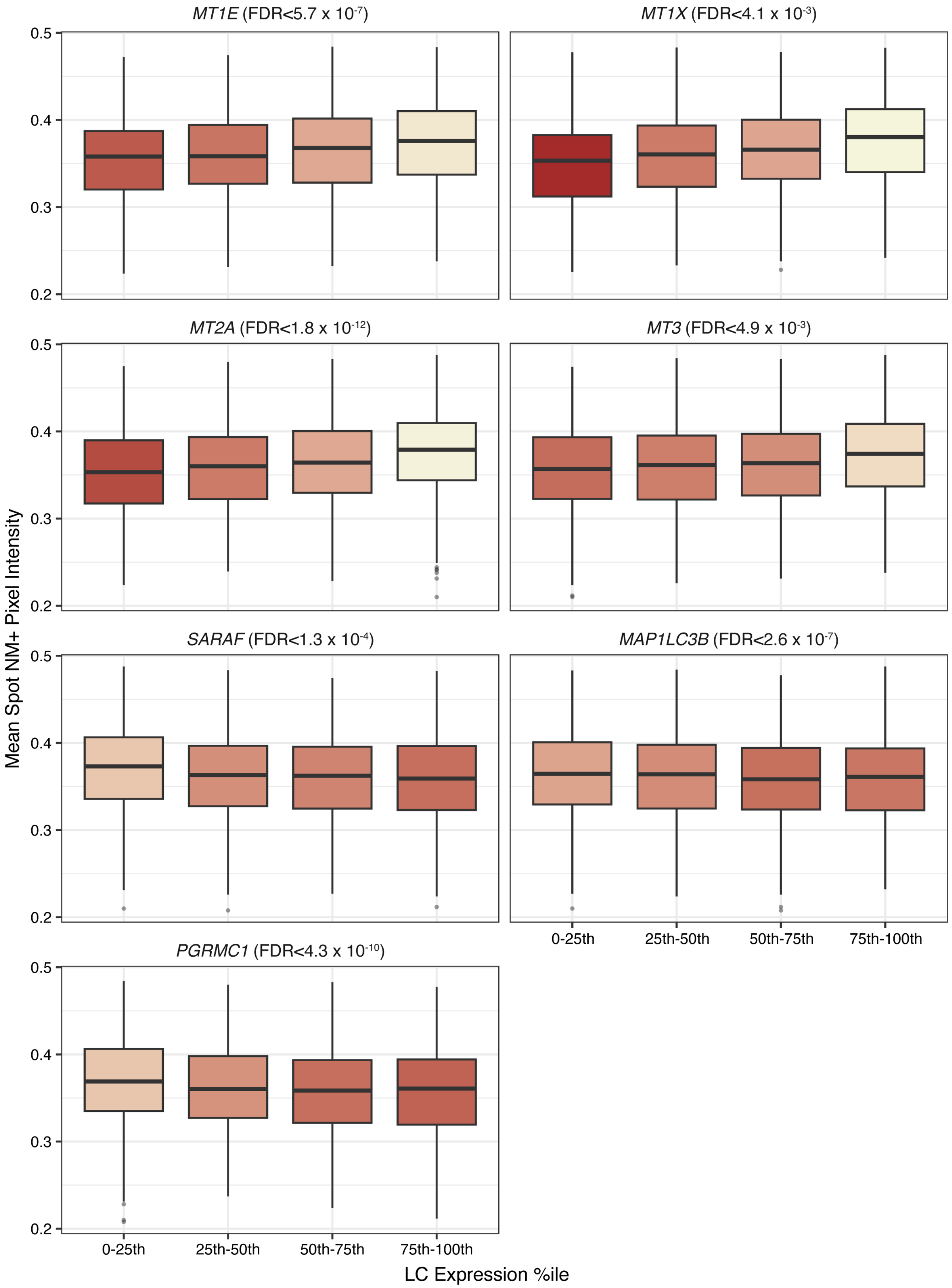


**Supplemental Figure S19. Metallothioneins and additional genes of interest associated with NM intensity**. For each gene, spot-wise mean NM pixel intensities within $LC^{NM+}$ are plotted against spot-level expression quartiles (excluding zero expression spots) in all samples containing ≥20 $LC^{NM+}$ spots. Lower mean NM intensities signify darker NM.
